# Deep learning and CRISPR-Cas13d ortholog discovery for optimized RNA targeting

**DOI:** 10.1101/2021.09.14.460134

**Authors:** Jingyi Wei, Peter Lotfy, Kian Faizi, Sara Baungaard, Emily Gibson, Eleanor Wang, Hannah Slabodkin, Emily Kinnaman, Sita Chandrasekaran, Hugo Kitano, Matthew G. Durrant, Connor V. Duffy, Patrick D. Hsu, Silvana Konermann

## Abstract

Transcriptome engineering technologies that can effectively and precisely perturb mammalian RNAs are needed to accelerate biological discovery and RNA therapeutics. However, the broad utility of programmable CRISPR-Cas13 ribonucleases has been hampered by an incomplete understanding of the design rules governing guide RNA activity as well as cellular toxicity resulting from off-target or collateral RNA cleavage. Here, we sought to characterize and develop Cas13d systems for efficient and specific RNA knockdown with low cellular toxicity in human cells. We first quantified the performance of over 127,000 RfxCas13d (CasRx) guide RNAs in the largest-scale screen to date and systematically evaluated three linear, two ensemble, and two deep learning models to build a guide efficiency prediction algorithm validated across multiple human cell types in orthogonal validation experiments (https://www.RNAtargeting.org). Deep learning model interpretation revealed specific sequence motifs at spacer position 15-24 along with favored secondary features for highly efficient guides. We next identified 46 novel Cas13d orthologs through metagenomic mining for activity and cytotoxicity screening, discovering that the metagenome-derived DjCas13d ortholog achieves low cellular toxicity and high transcriptome-wide specificity when deployed against high abundance transcripts or in sensitive cell types, including human embryonic stem cells, neural progenitor cells, and neurons. Finally, our Cas13d guide efficiency model successfully generalized to DjCas13d, highlighting the utility of a comprehensive approach combining machine learning with ortholog discovery to advance RNA targeting in human cells.

## Introduction

The ability to perturb desired RNA molecules with high efficiency and specificity is required for functional elucidation of the transcriptome and its diverse phenotypes. Despite rapid progress in effective technologies for genome engineering, analogous systems for transcriptome engineering lag behind their DNA counterparts. While RNAi has long been used for RNA knockdown, it is challenging to engineer and suffers from widespread off-target effects (Jackson et al., 2003; Sigoillot et al., 2012) due to its important role in endogenous miRNA processing (Doench et al., 2003). The discovery and development of RNA-guided RNA-targeting CRISPR systems, such as Cas13 enzymes, provides an orthogonal and modular approach to overcome these limitations (Abudayyeh et al., 2016; East-Seletsky et al., 2016). Because CRISPR proteins are orthogonal to eukaryotic systems, they can be easily engineered to bind or cleave target RNA molecules. Further, their modular nature enables the facile fusion of effector domains to expand effector functionality. As a result, a broad suite of Cas13-based tools is now able to perturb RNA expression (Abudayyeh et al., 2017; Konermann et al., 2018) or splicing (Konermann et al., 2018), mediate RNA editing (Abudayyeh et al., 2019; Cox et al., 2017; Xu et al., 2021) or methylation (Wilson et al., 2020), as well as profile RNA-protein interactions (Han et al., 2020). These capabilities are now accelerating applications across the study of fundamental RNA biology, RNA-based therapeutics, and molecular diagnostics.

The Cas13 family is unified by the presence of two conserved HEPN ribonuclease motifs, and these enzymes are activated by binding to cognate target RNA as specified by the Cas13 guide RNA (Abudayyeh et al., 2016; East-Seletsky et al., 2016; Slaymaker et al., 2021; Zhang et al., 2018). Several subtypes have been defined on the basis of sequence diversity and domain architecture. Cas13d enzymes – in particular the engineered Cas13d from *R. flavefaciens* strain *XPD3002* (CasRx) (Konermann et al., 2018) – are the smallest and most efficient Cas13 RNA targeting effectors in mammalian and plant cells reported to date (Wessels et al., 2020; Li et al., 2021; Mahas et al., 2019), motivating their further characterization and optimization as RNA targeting tools. In order to successfully apply Cas13d in high-throughput applications, the ability to design highly effective guide RNAs is critical. Recent efforts to understand and predict Cas13d guide activity have taken a first step in this direction, by using a dataset of 2,918 guide RNAs across four transcripts to train a random forest model (Wessels et al., 2020) and by using combined datasets of 10,279 guides to train a deep learning model (Cheng et al. 2023). In addition to the relatively small datasets, the manual selection of guide sequence features (Wessels et al., 2020) or lack of secondary features (Cheng et al. 2023) has limited a broader understanding of Cas13d targeting preferences.

Here, we conducted the largest Cas13d screen to date, quantifying CasRx guide efficiency across >127,000 guide RNAs tiling 55 essential transcripts by measuring their effects on cell proliferation in human cells. We systematically compared a series of computational models on this dataset to predict guide activity. A deep learning convolutional neural network (CNN) model was able to most accurately predict highly effective guides. Model interpretation enabled us to discover a preferred sequence motif at spacer position 15-24 along with a preference for low guide free energy and high target region accessibility for high efficiency guides. We validated the model against orthogonal datasets and confirmed high accuracy across target transcripts and five different cell types.

Across the Cas13 subtypes structurally characterized to date, the RNA cleavage site formed by the two HEPN domains is located distal to the guide binding groove (Liu et al., 2017; Slaymaker et al., 2021; Zhang et al., 2018), which can result in the cleavage of non-target bystander RNA molecules (known as ‘collateral’ cleavage) *in vitro* by the HEPN domains activated upon target RNA binding. Initial reports for Cas13a, b, and d systems in routinely used mammalian cell lines reported a low degree of off-target effects in eukaryotic cells (Abudayyeh et al., 2017; Cox et al., 2017; Konermann et al., 2018). However, more recently, several groups reported cellular toxicity and more pronounced off-target effects of CasRx, LwaCas13a, and PspCas13b in sensitive cell types (Ai et al., 2022; Özcan et al., 2021), *in vivo* (Buchman et al., 2020), and when targeting highly expressed transcripts (Shi et al. 2023).

To understand if this cellular toxicity is shared across Cas13 orthologs, we computationally identified 46 novel Cas13d orthologs from recently reported prokaryotic genomes and metagenomic contigs and screened them for target transcript knockdown activity and cytotoxic effects in human cells. We identified DjCas13d, a highly efficient ortholog with minimal detectable cellular toxicity when targeting highly expressed transcripts across multiple cell types, including human embryonic stem cells, neural progenitor cells, and neurons.

Furthermore, we show that our CasRx-based guide design model extends to DjCas13d and accurately selects highly efficient guides, illustrating its generalizability across effectors and cell types. Overall, we advance the transcriptome engineering toolbox by developing a robust Cas13d guide design algorithm based on a high-throughput guide screen (https://www.RNAtargeting.org), and identifying a compact and high-fidelity Cas13d ortholog for efficient RNA targeting. Finally, we outline a strategy to systematically develop and interpret robust deep learning models for sequence-based classification.

## Results

### Deep learning of Cas13d guide RNA efficiency based on large-scale transcript essentiality screening

In order to systematically understand factors impacting Cas13d guide efficiency, we generated a library of more than 100,000 RfxCas13d (CasRx) guide RNAs and evaluated their efficiency in a large-scale pooled screen. Reasoning that CasRx knockdown of essential transcripts would lead to the depletion of highly effective guides due to reduced cellular proliferation, we selected a set of 55 essential genes identified in three previously reported survival screens performed with RNAi and CRISPR interference (CRISPRi) in K562 cells (Hart et al., 2015; Horlbeck et al., 2016; Luo et al., 2008) for a proliferation-based survival screen. K562 cells were selected due to their ease of use in pooled screens and our observation of variable CasRx-mediated endogenous protein knockdown in this cell line **(Figures S1A, B).**

To perform the screen, we first generated stable K562 cell lines via transfection of an all-in-one plasmid encoding the CasRx effector, PiggyBac transposase, and an antibiotic selection cassette. Next, we designed CasRx guides that tile the 5’ UTR, coding sequence (CDS), and 3’ UTR of the 55 essential transcripts with single nucleotide resolution. As controls, we designed guides tiling 5 non-essential transcripts as well as 3,563 non-targeting guides. The effector cell line stably expressing CasRx was transduced with a pooled lentiviral library containing all 144,745 guide RNAs. Cells were cultured for 14 days, after which we analyzed guide abundances by NGS and computed a depletion ratio for each guide compared to its original abundance in the input library (**Figure 1A)**. Analysis of the cumulative distribution of guide d14/input ratio demonstrated that the top 20th percentile of guides targeting essential transcripts are clearly separated from guides targeting non-essential transcripts or non-targeting guides **(Figure 1B)**.

**Figure 1:**
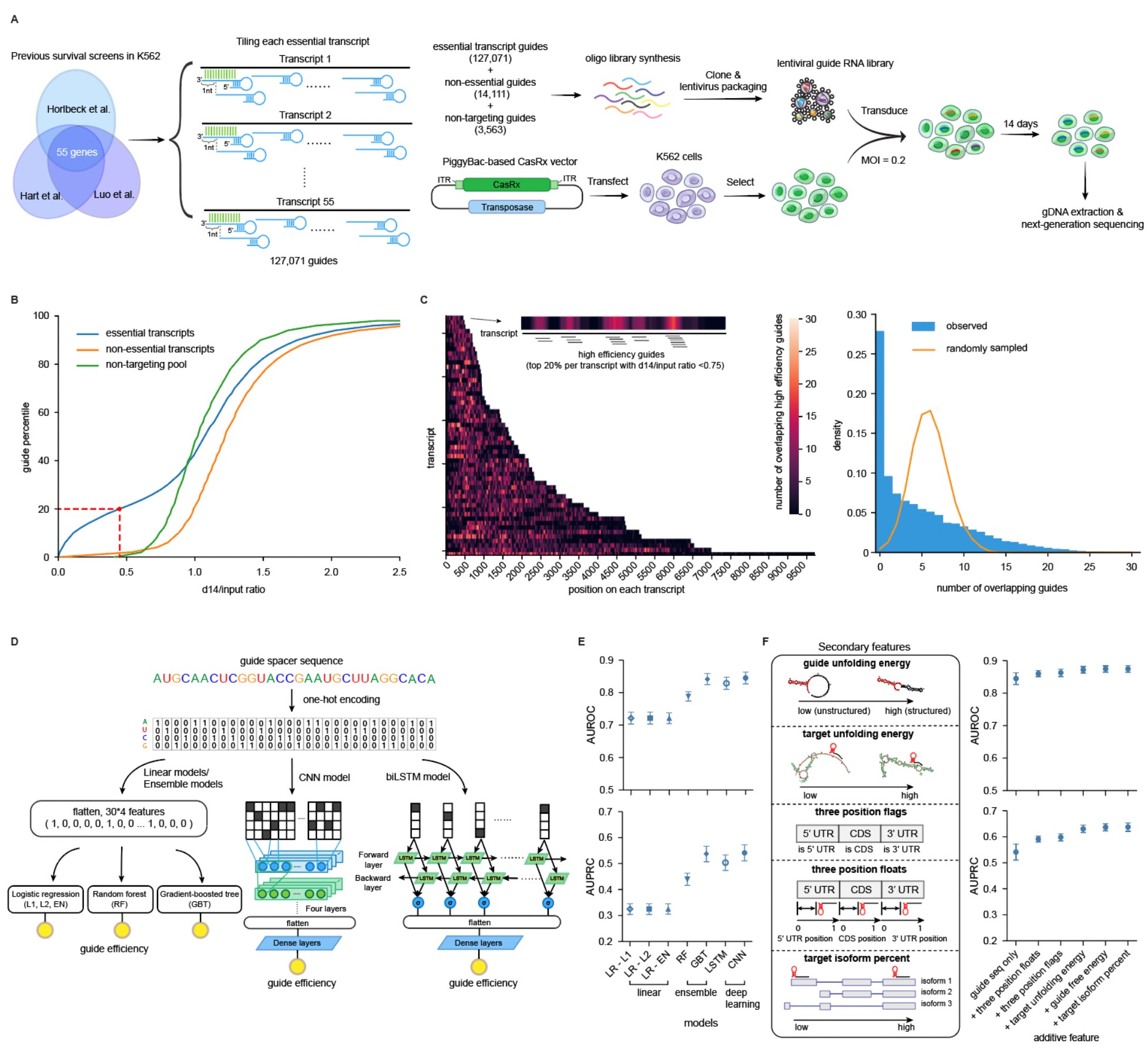
Deep learning of Cas13d guide RNA efficiency based on large-scale transcript essentiality screening. **A.** Schematic of the pooled CasRx guide tiling screen for essential transcript knockdown as a readout of per-guide knockdown efficiency. Over 127,000 targeting guide RNAs were included. **B.** Cumulative distribution of the ratio of relative guide abundance at day 14 compared to the input library across guides targeting essential gene transcripts (blue), non-essential gene transcripts (orange), and non-targeting guides (green). The red dashed line indicates the ratio at the top 20th percentile of essential transcript targeting guides. **C**. Heat map of the positional distribution of high efficiency guides along each transcript. From here forward, high efficiency guides are defined as the top 20% guides within each transcript with a d14/input ratio lower than 0.75 after essential off-target filtering. Heat map color indicates the number of overlapping high efficiency guides at each nucleotide position along the transcript, and the histogram (right panel) depicts the observed frequency distribution of these data (blue) as compared to a random distribution of 20% of guides in the library (orange curve). **D.** Schematic of the computational algorithms assessed in this study to predict guide efficiency based on spacer sequence alone. **E.** Comparison of prediction accuracy between linear, ensemble and deep learning models across 9-fold splits of held-out transcripts. Averages of Area Under the Receiver Operating Characteristic curve (AUROC) and Area Under the Precision-Recall Curve (AUPRC) across test sets from all 9 folds are shown ± SD. LR - L1, logistic regression with L1 regularization (Lasso Regression); LR - L2, logistic regression with L2 regularization (Ridge regression); LR - EN, logistic regression with elastic net regularization (Elastic Nets) ; GBT, Gradient-Boosted Tree; RF, Random Forest classifier; CNN, Convolutional Neural Network; biLSTM, Bidirectional long short-term memory neural network. Note that the baseline for AUPRC is equal to the fraction of positive class (high efficiency guides), in this case 0.18. **F.** Secondary features were evaluated for their ability to improve sequence-only model performance. Each secondary feature (or feature group) was added to the CNN model sequentially, ordered by its individual contribution to model performance in **Figure S3G**. AUROC and AUPRC (mean ± SD) of all test sets from the 9-fold split of transcripts are shown.

Essential transcripts may vary in their magnitude of impact on cell proliferation and survival upon depletion. A transcript-level analysis of guide depletion confirmed this expectation (**Figure S1C**). In order to compensate for this in our analysis going forward, we selected the most effective guides for each individual transcript (see Methods for a full description of selection parameters) as high efficiency guides. A heat map representation of the positions of these high efficiency guides within each target transcript revealed a striking degree of clustering, leading to guide hot spots and deserts along the transcript and clearly deviating from a random distribution **(Figure 1C**). Multiple factors could be responsible for the observed clustering of high efficiency guides, including sequence-, structure-, or position-based effects of the guide RNA or target transcript.

### Prediction of CasRx guide activity based on guide RNA sequence alone

We sought to systematically analyze these potential features that could distinguish high efficiency Cas13d guides and develop computational algorithms to predict guide efficiency. Initial analysis of the correlation of nucleotide identity with guide efficiency at each position along the 30 nt spacer showed a preference for G and C at the direct repeat-proximal spacer positions 15-24 (**Figure S2A**). Therefore, we reasoned that spacer sequence alone might be predictive of guide efficiency when used as model input. We then developed a series of computational models for prediction of guide efficiency based on one-hot encoding of the 30 nt guide spacer sequence without manual sequence feature selection. To understand the impact of computational model type, we systematically built and assessed the following models: 3 linear models employing logistic regression (Lasso Regression (L1), Ridge regression (L2) or Elastic Net (EN)), 2 ensemble models (Random forest (RF) and Gradient-boosted tree (GBT)) and 2 deep learning models (convolutional neural network (CNN) and bidirectional long short-term memory neural network (LSTM)) (**Figure 1D**).

All of these models were trained to classify high efficiency guides for target transcripts. Due to the observed high degree of clustering of effective guides along a transcript (**Figure 1C**), models that are tested on held-out guides from the same transcripts they were trained on would potentially be subject to overfitting by learning the targeting hotspots specific to those transcripts. To alleviate overfitting and ensure model generalizability to other transcripts, we employed 9-fold cross-validation on the 54 target transcripts (leaving out *RPS19BP1* as it clustered with non-essential transcripts (**Figure S1C**)), with models being trained and tested on non-overlapping sets of transcripts. We compared the performance of all 7 models and observed high model performance for the gradient-boosting tree (GBT) and the two deep learning models based on Area Under the Receiver Operating Characteristic curve (AUROC), which evaluates prediction accuracy for both the positive class (high efficiency guides) and the negative class, and Area under the Precision-Recall Curve (AUPRC) metrics, which focuses primarily on the prediction accuracy of the positive class (high efficiency guides), across all 9 fold splits (**Figure 1E**).

Overall, the CNN model performed best with a high AUROC of 0.845 (relative to a baseline of 0.5) and a high AUPRC of 0.541 (relative to a baseline of 0.18), so we chose this model for further refinement and evaluation. The high prediction accuracy of this model based on the spacer sequence alone indicates that sequence is a primary factor determining guide efficiency. We further determined that the addition of target flanking sequences of varying length from 1-7 nt to the CNN model did not meaningfully improve model performance (**Figure S2B**), consistent with our previous biochemical studies suggesting a lack of strong flanking sequence requirements (Konermann et al., 2018). To understand the minimal spacer length required for accurate prediction, we computationally truncated the spacer sequence from the 3’ end in the CNN model input, and found only a minor impact on model accuracy until reaching a spacer length of 24 nt, after which a gradual drop in AUROC and AUPRC was observed (**Figure S2C**). We validated this experimentally, demonstrating decreasing target knockdown when using guides shorter than 24 nt in spacer length (**Figure S2D**).

### Addition of secondary features improves guide efficiency prediction accuracy

Beyond guide sequence alone, secondary guide attributes such as guide unfolding energy or target site position (CDS or UTR) may impact guide performance. To understand their potential contribution, we first evaluated the correlation of such secondary features with guide efficiency (**Figure 1F schematics, S3A-F**). We found that higher predicted guide and target RNA unfolding energy, implying more highly structured RNA sequences, were predictive of poor guide efficiency. We also observed a preference for intermediate spacer GC content (45-55%), guides targeting the coding region (CDS), as well as guides targeting regions conserved across transcript isoforms.

As most of the secondary features investigated exhibited a modest correlation with guide efficiency, we tested whether they would improve model performance when added to the spacer sequence-only CNN model. When adding these features individually, we found that the guide target site position had the most prominent effect, followed by target and guide RNA folding energy (**Figure S3G**). The addition of spacer GC content did not significantly improve model performance, consistent with our expectation that this feature has been successfully captured by the spacer sequence-only CNN model. Sequentially including each secondary feature ranked by their individual contribution into the sequence-only CNN model, we found that AUROC and AUPRC were improved with each addition, leading to a final model with a very high average AUROC of 0.875 and a high average AUPRC of 0.638 (**Figure 1F** and **S3H-J** for feature variations). Adding the same set of secondary features also improved the GBT model (**Figure S4**), the best performing model not based on deep learning, indicating the contribution of these secondary features to guide efficiency.

One of the key applications of a predictive model like this one would be to accurately predict the most effective guides in order to aid in guide and library design. The CNN model returns a float score ranging from 0 to 1 for every guide, and different thresholds can be chosen for high efficiency guide classification. To evaluate model performance for optimal guide selection, we set a high model score threshold of 0.8 and plotted the true percentile rank distribution of the guides above the score threshold. As expected, the guides were heavily skewed towards the highest efficiency ranks, with a true positive ratio of 0.83 (83% being true high efficiency guides (top 20th percentile)). Setting an even more stringent model score threshold to 0.9 further increased the true positive ratio to 93% (**Figure S3K**).

### Model interpretation reveals favored sequence and secondary features of high efficiency guides

Having built high performance models that accurately predict efficient guides, we asked whether these models could help us understand the features contributing to guide efficiency by using three model interpretation methods. We first used an integrated gradients approach (IG) (Sundararajan et al., 2017) to provide observability for our CNN model. We began with the guide sequence preferences learned by the model, and IG analysis on each position in the guide spacer sequence nominated a core region of position 15-24 as a major contributor to guide efficiency (**Figure 2A**). Consistent with our original correlation analysis (**Figure S2A**), IG analysis on each positional nucleotide in the guide sequence revealed a clear preference for an alternating stretch of guanines, cytosines and guanines (G_15-18_C_19-22_G_23-24_) in this core region (**Figure 2B**).

**Figure 2:**
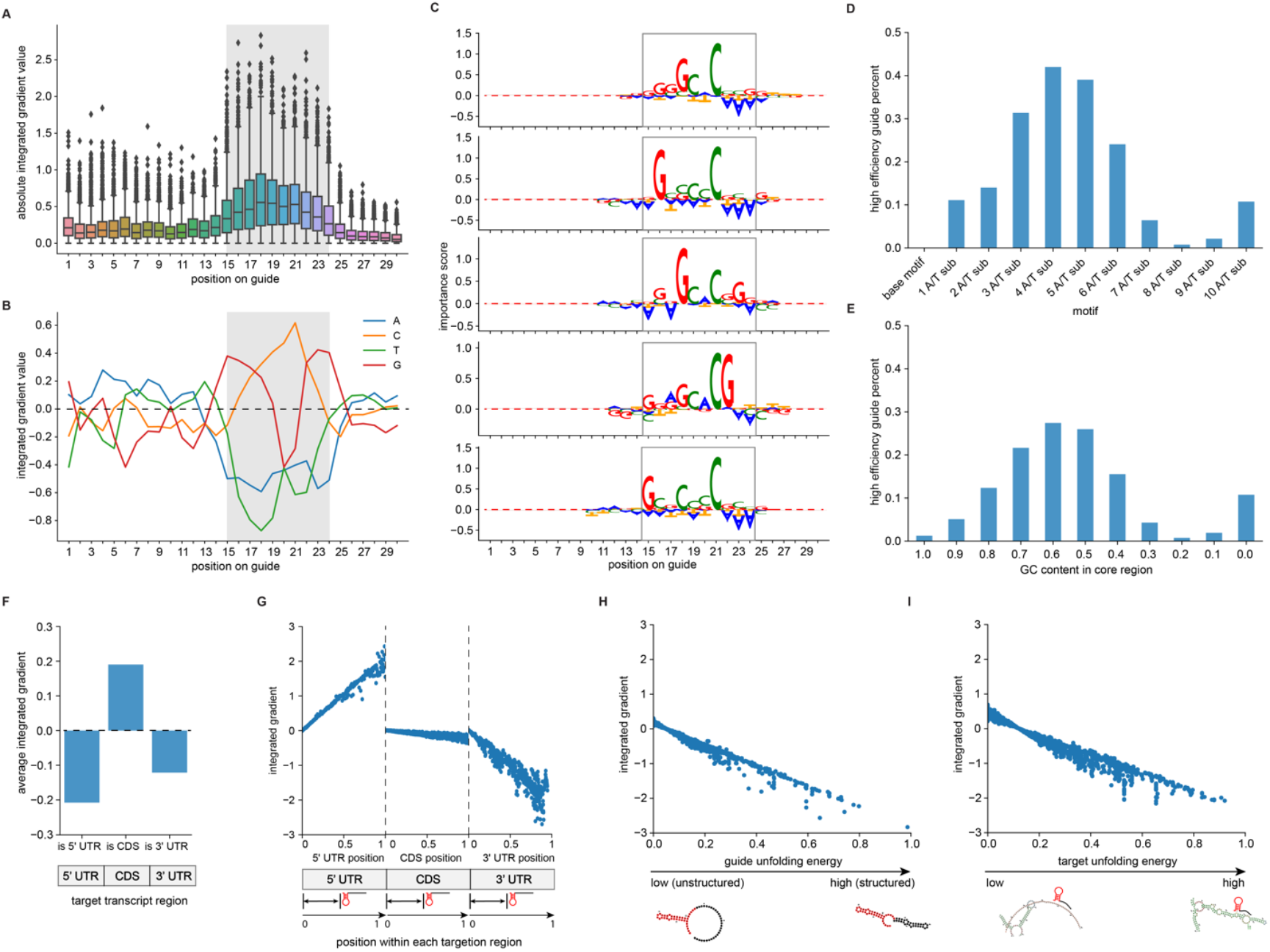
Deep learning model interpretation reveals favored sequence motifs and secondary features of high efficiency guides. **A.** Evaluation of the importance of each position in the guide spacer sequence in the CNN model using Integrated Gradients (IG). Higher absolute gradient values indicate greater importance for predicting a high efficiency guide. The gray box highlights the identified core region (position 15-24). **B.** Evaluation of the importance of each positional nucleotide in the guide sequence in the CNN model by IG. **C.** Top 5 sequence patterns identified by TF-MoDISco (Transcription Factor Motif Discovery from Importance Scores) in the CNN model. Patterns are aligned to the 30 nt spacer according to the mode position of the seqlets (sequence regions with high importance based on IG scores) in each pattern (**Figure S7A**). **D.** Fraction of high efficiency guides that contain the 10-base motif shown in panel B and A/T substitutions within the 10-base motif. **E.** Fraction of high efficiency guides across different core region GC content. Guides were divided into eleven bins based on the GC content in their core region (position 15-24), and the fraction of high efficiency guides belonging to each bin is plotted. **F.** Contribution of target transcript region (5’ UTR, CDS, or 3’UTR) to guide efficiency in the CNN model. The bar plots indicate average IGs of all test samples with different target position flags. **G.** Contribution of position within each transcript target region to guide efficiency in the CNN model. The scatter plots indicate individual IG values against individual input values across all test samples. The reference points are set to 0 for each transcript region. **H.** Contribution of predicted guide unfolding energy to guide efficiency in the CNN model. The reference point is set to 0. **I.** Contribution of predicted target unfolding energy to guide efficiency in the CNN model. The reference point is set to 0.

To confirm the favored sequence features across models and model interpretation methods, we further applied SHapley Additive exPlanations (SHAP), a game theoretic approach (Lundberg et al., 2020) to our GBT model, and a similar sequence preference in the same core region was observed (**Figures S5A, B**). In contrast, this unique sequence preference was not found for Cas13a when we performed a correlational analysis of available datasets (Abudayyeh et al., 2017; Metsky et al., 2022) (**Figure S6**). Indeed, no consistent sequence preference or core region emerged across the Cas13a datasets analyzed, which could be due to intrinsic enzymatic properties of Cas13a or limitations in the size of available datasets.

As our IG and SHAP analyses investigated each position in the guide sequence independently, we further sought to determine the role of specific motifs (nucleotide combinations) in guide efficiency. We employed Transcription Factor Motif Discovery from Importance Scores (TF-MoDISco), an algorithm that identifies sequence patterns or motifs incorporated in deep learning models by clustering important sequence segments based on per-position importance scores (Shrikumar et al., 2018). We discovered a total of 14 distinct sequence patterns associated with high efficiency guides from the CNN model, with the top 5 patterns shown in **Figure 2C**. As TF-MoDISco was initially applied for the identification of transcription factor binding motifs, it is designed to identify motifs in a position-independent manner. In our analysis, we noticed that all identified patterns were anchored to a specific position centered around guide spacer nucleotides 18-20 (**Figure S7A**), consistent with our prior observation of a core region.

Strikingly, all top 5 sequence patterns contained a cytosine at position 21, with a single guanine at varying positions in the core region across the different patterns. Taken together, the identified motifs can be summarized as **GN_x_C_21_** or **N_x_C_21_G** within the core region. Generally, the patterns were sparse and characterized by just two dominant bases (one G and one C), in contrast to the longer 10-base motif that the individual position-level analysis would have suggested (**Figures 2B and S5B**). Consistent with our results above, an analysis of enriched and depleted 3-mers in high efficiency guides across the spacer sequence revealed that enriched 3-mers were again clustered in the core region (position 15-24) (**Figure S7B**). In addition to the consistent finding of a prominent enrichment of C at position 21, they revealed a preference for A or T intercalated with G and C (**Figures S7B, C**), a finding that was obscured in the per-position analysis. Analysis of enriched and depleted 4-mers in high efficiency guides also led to a similar finding (**Figure S7D**). A/T substitutions within the 10-base motif (G_15-18_C_19-22_G_23-24_) (**Figure 2D**) and analysis of the GC content in the core region (**Figure 2E**) for high efficiency guides further confirmed a preference for a medium GC content via A/T nucleotides at the N positions of the key **GN_x_C_21_** or **N_x_C_21_G** motif.

Next, we used IG and SHAP to investigate the contribution of secondary features in the CNN and GBT models. IGs revealed that targeting the beginning of the 5′ UTR and the end of the 3′ UTR was the most disfavored, while targeting the coding region (CDS) was generally favored, with a slight preference for the beginning of the CDS (**Figures 2F, G**). In agreement with our correlation analysis, guide and target unfolding energy also had a relatively high impact on guide efficiency, with lower unfolding energy favored by high efficiency guides (**Figures 2H, I**). SHAP analysis on our GBT model showed a consistent direction of feature contribution to guide efficiency (**Figure S5C**) and ranked spacer sequence composition as the most important feature.

Taken together, our systematic model interpretation was consistent across models and analysis approaches, was able to rank features by their contribution toward guide classification, and significantly expanded our understanding of preferred longer-range sequence motifs that were missed by simpler correlational analyses.

### Systematic validation of the guide efficiency model across 5 cell types with endogenous protein knockdown

Next, we sought to experimentally validate our model through CasRx-mediated knockdown of cell surface markers, reasoning that an orthogonal readout to transcript essentiality and cell survival would ensure generalizability of our model predictions to multiple readout modalities. To this end, we performed a screen using a library of 3,218 guides tiling the transcripts of two cell surface markers, *CD58* and *CD81*, with single-nucleotide resolution. 10 days after lentiviral transduction of the guide library, cells were FACS sorted into 4 bins on the basis of target protein expression level (**Figure 3A**) and the enrichment of individual guides in the top and bottom bins (exhibiting the greatest or least magnitude of knockdown, respectively) was assessed. We observed clear separation of the most efficient targeting guides from the non-targeting guides based on the enrichment ratio, with zero non-targeting guides appearing in the top 20th percentile of guide efficiency (**Figure 3B**).

**Figure 3:**
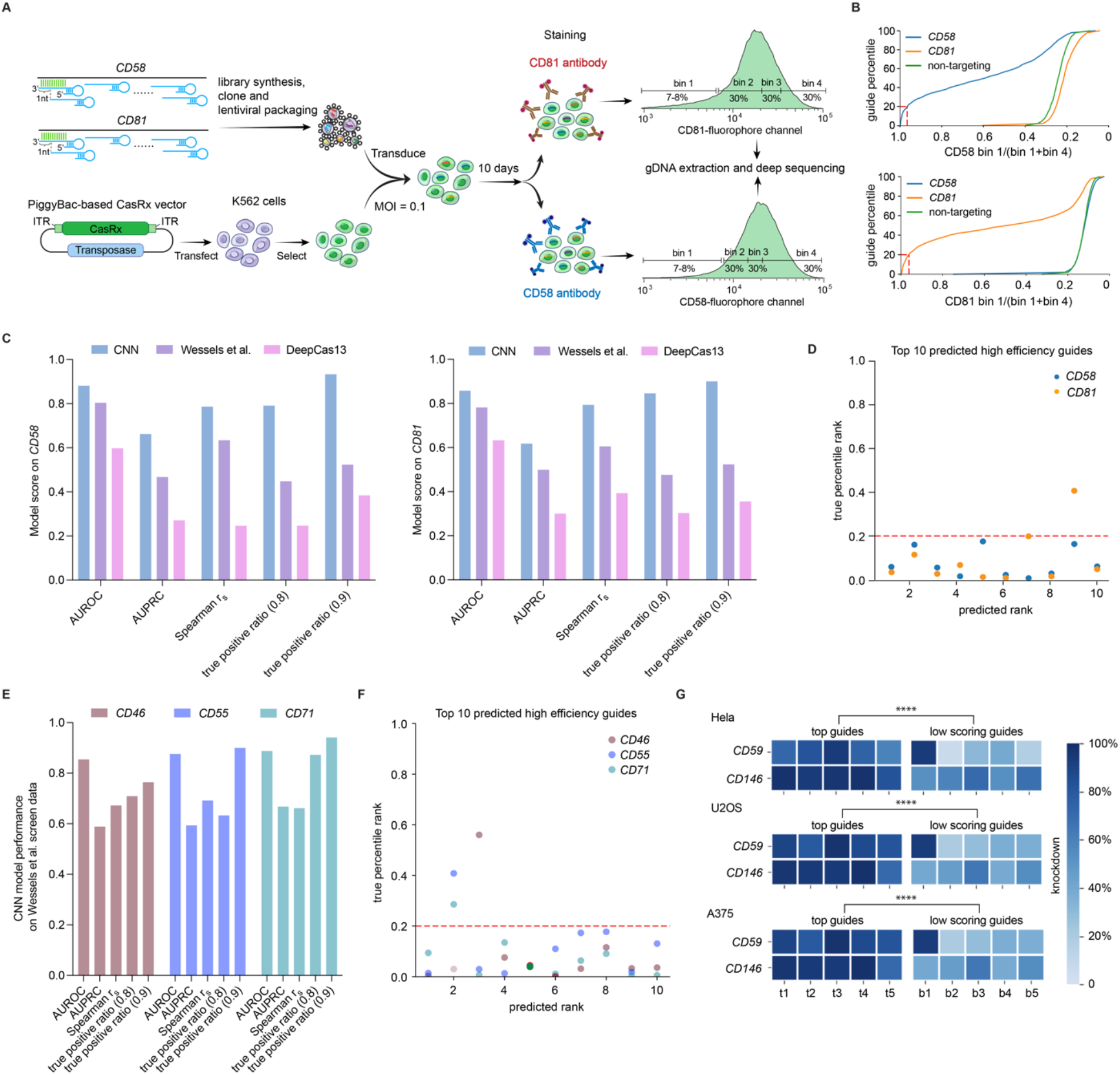
Systematic validation of the guide efficiency model across 5 cell types with endogenous protein knockdown. **A.** Schematic of the pooled CasRx guide tiling screen targeting *CD58* or *CD81* transcripts in K562 cells followed by flow cytometry-based readout of cell-surface CD58 or CD81 protein abundance. **B.** Cumulative distribution of guide enrichment ratios for *CD58*, *CD81* and non-targeting guide categories, calculated as the ratio of guide percentage in bin 1 (greatest knockdown) relative to the sum in bin 1 and bin 4 (least knockdown). Red dashed lines indicate the ratio for the top 20th percentile of targeting guides. **C.** Model comparison on *CD58* and *CD81* guides. CNN, the ensemble CNN model built on the survival screen data in this work; Wessels et al. model, a previously published CasRx random forest model (Wessels et al., 2020); DeepCas13, a previously published CasRx deep learning model (Cheng et al. 2023). Model performance is evaluated by AUROC, AUPRC, Spearman’s correlation coefficient (*r*_s_) and true positive ratio at 0.8 and 0.9 model score cutoffs across guides targeting *CD58* (left panel) and *CD81* (right panel). **D.** True percentile rank of the top 10 predicted high efficiency guides for *CD58* and *CD81*. The red dashed line indicates the top 20th percentile of *CD58-* or *CD81*-targeting guides. **E.** Performance of the ensemble CNN model on a published CasRx guide tiling dataset of three CD transcripts (*CD46*, *CD55*, and *CD71*) in HEK293FT cells (Wessels et al., 2020). Model AUROC, AUPRC, Spearman’s correlation coefficient (*r*_s_), and true positive ratio at 0.8 and 0.9 model score cutoffs are shown for each transcript. **F.** True percentile rank of the top 10 predicted guides by our model for three transcripts in a published CasRx guide tiling dataset in HEK293FT cells (Wessels et al., 2020) predicted by the ensemble CNN model. The red dashed lines indicate the top 20th percentile of targeting guides. **G.** Knockdown efficiency of the predicted 5 top scoring guides and 5 low scoring guides for two transcripts (*CD59* and *CD146*) measured by flow cytometry in Hela, U2OS, and A375 cells. Heat map color indicates the mean knockdown efficiency for each guide across n = 3 biological replicates. The top scoring guides and low scoring guides were significantly different at P<0.0001 for Hela, U2OS and A375 cells based on Welch’s t test.

We evaluated our CNN model’s performance on this new dataset and found that an ensemble CNN model comprising all 9 fold splits of the survival screen outperformed each individual split model (**Figure S8A**) and achieved high prediction accuracy for both *CD58* (AUROC of 0.88 and AUPRC of 0.66) and *CD81* (AUROC of 0.86 and AUPRC of 0.62) (**Figure 3C**). This performance is comparable to the model accuracy on held-out essential transcripts from our initial screen (**Figure 1F**), highlighting its generalizability. Compared with two existing Cas13d guide design models (Wessels et al., 2020, Cheng et al. 2023), our model showed the highest AUROC, AUPRC, and Spearman correlation. Importantly, we showed that at a 0.9 score cutoff, our model exhibited a very high true positive ratio of 0.93 and 0.9 for *CD58* and *CD81*, respectively, in contrast to the Wessels et al. model (0.52 for both *CD58* and *CD81*) and DeepCas13 (0.38 for *CD58* and 0.35 for *CD81*) (**Figure 3C**). The far higher true positive ratio at high score cutoffs underlines the superior utility of our model for key applications such as predicting the top 3-10 guides per target transcript in individual targeting or library-based screening approaches. Illustrating this use case, we examined the true percentile rank of the top 10 predicted high efficiency guides for *CD58* and *CD81*, showing that 10/10 guides for *CD58* and 9/10 for *CD81* were highly effective (**Figure 3D**).

To assess generalizability to other cell types, we evaluated our model’s performance on a published CasRx guide tiling dataset (∼3000 guides in HEK293FT cells from the Wessels et al. training dataset). Our model showed high AUROC (0.85, 0.88 and 0.85 for *CD46, CD55 and CD71,* respectively), AUPRC (0.59, 0.59 and 0.67), Spearman correlation (0.67, 0.69 and 0.66), and true positive ratio (0.76, 0.9 and 0.94 at a 0.9 score cutoff) (**Figure 3E**). Among the top 10 predicted high efficiency guides, 90% were highly efficient (falling into the top 20% percentile of efficient guides) (**Figure 3F**). When compared against the Wessels et al. model on opposing datasets (**Figure S8B)**, our model showed significantly higher prediction accuracy using all evaluation metrics (AUPRC: 0.617 vs 0.379; Spearman correlation rs: 0.675 vs 0.391; AUROC: 0.873 vs 0.733; true positive ratio (0.9 cutoff): 87% vs 51%), further supporting the generalizability and high performance of our model.

As a final test of the ability of our model to predict efficient guides for knockdown of desired transcripts in different cell types, we selected 5 top scoring guides and 5 low scoring guides (excluding the very bottom of our ranking) for two different transcripts (*CD59* and *CD146*), and tested the knockdown efficiency of each guide in Hela, U2OS, and A375 cells (**Figure 3G**). Across all three cell lines, the top scoring guides showed very efficient target knockdown (72%-98% with a median of 90%) while low scoring guides showed variable and significantly lower levels of knockdown (6%-70% with a median of 35%), confirming the utility and generalizability of our model across 5 cell types (K562, HEK293FT, Hela, U2OS, and A375).

### Discovery of DjCas13d, a high-efficiency RNA targeting enzyme with minimal cellular toxicity in human cells

In genome engineering, two of the most important features are efficiency and specificity. A key emerging limitation of several Cas13 systems is the induction, in certain contexts, of cellular toxicity by its RNA trans-cleavage activity (Ai et al., 2022; Buchman et al., 2020; Özcan et al., 2021), hampering their application as a generalizable transcriptome engineering tool. In the context of this study, we also observed various degrees of cellular toxicity for CasRx when paired with highly efficient guides in the A375 cell line (**Figure S9A).**

To address this, we reasoned that the evolutionary diversity of Cas13d enzymes may have already developed solutions to these challenges. To develop a more broadly useful transcriptome engineering tool, we sought to identify a Cas13d ortholog that combines the key positive traits of CasRx, like its small size and high targeting efficiency, with low cellular toxicity. We applied our previously described computational approach for Cas13d discovery (Konermann et al., 2018) to an expanded database of metagenomic datasets and discovered 46 previously uncharacterized Cas13d enzymes, expanding the known Cas13d family from 7 to 53 members (**Figure 4A**, **Table S7**).

**Figure 4:**
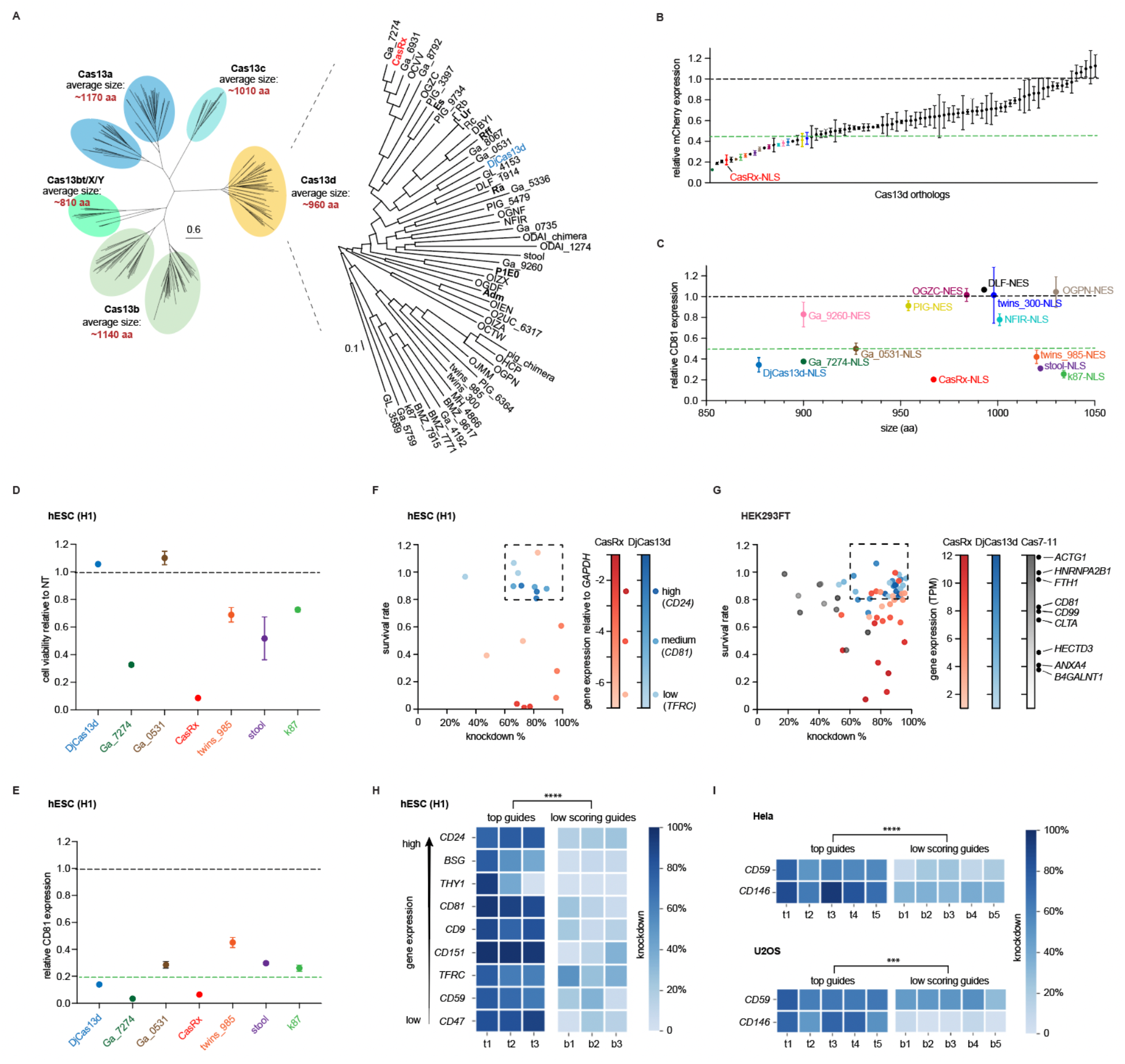
Discovery of DjCas13d, a high-efficiency RNA targeting enzyme with minimal cellular toxicity in human cells. **A.** Phylogenetic tree of Cas13 enzymes including the expanded Cas13d subtype clade (yellow). 46 additional Cas13d orthologs were identified through mining of recent metagenomic datasets. The 7 previously identified Cas13d orthologs including CasRx (red) are shown in bold text. The newly discovered ortholog DjCas13d is shown in blue. All the ortholog sequences are provided in Table S7. **B.** Evaluation of the knockdown efficiency of all Cas13d orthologs shown in panel A on an mCherry reporter transcript in HEK293FT. Horizontal green dashed line denotes our selected cutoff of >55% knockdown efficiency; the hits are color-coded for further study. **C.** Evaluation of the knockdown efficiency of the selected 14 Cas13d orthologs on an endogenous transcript, *CD81*, as measured by flow cytometry-based readout of protein abundance. The horizontal green dashed line denotes a 50% knockdown efficiency cutoff. Cas13d enzymes are plotted in order of their protein size on the x-axis (small to large). **D-E.** Evaluation of cell viability (panel **D**) and knockdown efficiency (panel **E**) of cells expressing each of the top seven most efficient Cas13d orthologs in H1 hESCs along with a *CD81-*targeting guide. The horizontal green dashed line in panel E denotes an 80% knockdown efficiency cutoff. Orthologs are ordered by their size and color-coded as in panel C. Values are shown as mean ± SEM for n = 3 replicates. **F.** Evaluation of cellular viability (y axis) and knockdown efficiency (x axis) of DjCas13d and CasRx across three transcripts in the hESC line, H1. Three top guides were picked for each transcript based on the CNN model score. Each dot on the scatter plot represents one guide’s survival rate and knockdown (mean for n = 3 replicates). The dots are colored by the effector used (CasRx: red, DjCas13d: blue), and the color gradients denote the expression level of the target transcript relative to *GAPDH* (log2 relative expression) in the hESC line H1 based on qPCR. The dashed box denotes guides with >80% survival rate and >60% knockdown. 89% of DjCas13d guides are within the box while only 11% of CasRx guides are within the box. **G.** Evaluation of cellular viability (y axis) and knockdown efficiency (x axis) of DjCas13d, CasRx, and Cas7-11 across nine transcripts of different expression levels in HEK293FT using the same spacer sequences across all three enzymes. Three top guides were picked for each transcript based on the CNN model score. Each dot on the scatter plot represents one guide’s survival rate and knockdown (mean for n = 3 replicates). The dots are colored by the effector used (CasRx: red, DjCas13d: blue, Cas7-11: grey), and the color gradients denote the expression level (TPMs (transcript per million), log2(TPM+1)) of the target transcript. As in panel F, the dashed box denotes guides with >80% survival rate and >60% knockdown. 84% of DjCas13d guides are within the box while 32% of CasRx guides and 0 Cas7-11 guides are within the box. **H.** Knockdown efficiency of DjCas13d paired with 3 top scoring guides and 3 low scoring guides from the CNN model prediction on nine transcripts of different expression levels in H1 hESCs. Heat map color indicates the mean extent of knockdown for each guide across n = 3 biological replicates. The top scoring guides and low scoring guides were significantly different at P<0.0001 based on Welch’s t test. **I.** Knockdown efficiency of DjCas13d paired with 5 top scoring guides and 5 low scoring guides from the CNN model prediction on two transcripts (*CD59* and *CD146*) in Hela and U2OS cells. Heat map color indicates the mean knockdown efficiency for each guide across n = 3 biological replicates. The sets of top scoring guides and low scoring guides were significantly different at P<0.0001 in Hela and P<0.001 in U2OS based on Welch’s t test.

To evaluate these novel Cas13d enzymes for mammalian transcript knockdown, we synthesized human codon-optimized constructs of each enzyme with NLS (nuclear localization sequence) and NES (nuclear export sequence) fusions and measured their ability to knockdown the mCherry reporter transcript using a matched guide array containing two mCherry targeting guides. We identified 14 enzymes exhibiting >55% knockdown efficiency (**Figure 4B**) in this reporter assay. Because reporter knockdown is often weakly predictive of Cas13 performance on endogenous targets, we further tested the 14 orthologs on our shortlist for their knockdown efficiency when targeted to the endogenous *CD81* transcript. With this more stringent test, 7 orthologs exhibited >50% knockdown efficiency (**Figure 4C**), and we focused on these for further characterization.

Having identified this shortlist of the most efficient Cas13d enzymes, we next evaluated their cytotoxic effects in human embryonic stem cells (hESC), since we previously observed issues in this cell type with CasRx. When targeting the non-essential transcript *CD81* in this highly sensitive cell type, we were able to observe a significant reduction in viable cells expressing CasRx and most of the other Cas13d orthologs (**Figure 4D**), consistent with cytotoxic effects on other sensitive cell types reported in the literature (Özcan et al., 2021). Strikingly, two of the orthologs we tested (DjCas13d and Ga_0531) led to no detectable reduction of viable cell counts (**Figure 4D**). Of those two, we chose DjCas13d for additional characterization given its high knockdown efficiency (>80% in hESCs) (**Figure 4E**) and unusually small size (877aa, compared to 967aa for CasRx) (**Figure 4C**).

In a further evaluation across three guides each for three transcripts in hESCs, DjCas13d showed no significant effects on viable cell counts in contrast to CasRx, which caused significantly reduced viable cell counts in eight out of nine guides (**Figure 4F**). In terms of knockdown efficiency, DjCas13d showed high knockdown efficiency of >70% for most guides tested (median of 71.5%) – efficiency that was comparable to CasRx (median of 77.4%) (**Figure 4F**).

### DjCas13d induces minimal cellular toxicity when targeting highly expressed transcripts

Recent work (Ai et al. 2022; Shi et al. 2023) and our results in stem cells (**Figure 4F**) highlighted high target transcript abundance as a key variable for Cas13-mediated cellular toxicity in addition to the importance of cell type. In our own experiments in hESCs, we also observed the lowest survival rate for CasRx when targeting the most abundant transcript – *CD24* – while no such impact was observed for DjCas13d (**Figure 4F**). In order to further compare CasRx and DjCas13d under conditions known to promote cellular toxicity, we targeted three previously described highly expressed transcripts (*ACTG1*, *HNRNPA2B1*, *FTH1*) (Shi et al. 2023) in HEK293FT cells and confirmed a significant reduction of the number of viable cells when using CasRx but not DjCas13d (**Figure 4G**, all guides significant at P<0.0001). We targeted three medium- and three low expression level transcripts, confirming that lower expression of the target transcript alleviated the toxicity induced by CasRx (**Figure 4G**), consistent with initial reports (Konermann et al., 2018). By contrast, we observed minimal impact on viable cell counts when using DjCas13d to target any of these transcripts (**Figure 4G**), despite comparable knockdown efficiency of DjCas13d (knockdown median of 88%) to CasRx (median of 84%).

In a second head-to-head comparison, we tested DjCas13d against the recently reported Cas7-11 enzyme, which does not belong to the Cas13 family of CRISPR enzymes and was reported to have no impact on cell viability due to its distinct RNA cleavage mechanism (Kato et al., 2022; Özcan et al., 2021). We demonstrate that both DjCas13d and Cas7-11 have a comparably low impact on cell viability and proliferation (90% median cell count for DjCas13d across all targeting conditions, and 73% for Cas7-11) when targeting the same medium to highly expressed transcripts - in stark contrast to CasRx (46% median cell count). However, Cas7-11 suffered from diminished knockdown efficiency (median of 57%) compared to DjCas13d and CasRx (median of 88% and 84%, respectively) (**Figure 4G**).

Overall, we conclude that DjCas13d combines the best features of CasRx and Cas7-11, exhibiting low cellular toxicity and high knockdown efficiency. 84% of guides tested with DjCas13d showed >80% survival rate and >60% knockdown, while only 32% of CasRx guides and no Cas7-11 guides met these cutoffs.

### DjCas13d activity can be accurately predicted with our guide efficiency model

Given that DjCas13d belongs to the same subtype of CRISPR effectors as CasRx, we next sought to test whether our Cas13d guide design model could be successfully applied to this new Cas13d ortholog. Encouragingly, our data in **Figure 4F** and **G** demonstrated high efficacy of knockdown with guides recommended by the model when using DjCas13d across 12 transcripts of different expression levels and in different cell types. To further explicitly validate the model performance for DjCas13d, we selected a set of top and bottom scoring guides for a total of eleven transcripts across a range of expression levels in hESCs, HeLa, and U2OS cell lines.

Across hESCs (**Figure 4H**) as well as Hela and U2OS cells (**Figure 4I**), the predicted high efficiency guides resulted in a significantly higher degree of protein knockdown (median of 73.9%) compared with low-scoring guides (median of 19.7%) (**Figures 4H, I**). Altogether, these results demonstrate that our model generalizes to the novel DjCas13d ortholog, resulting in reliable knockdown performance and lack of apparent cellular toxicity even in sensitive cell types and for highly abudant transcripts. Given that the sequence divergence between DjCas13d and CasRx (29.9%) is similar to the divergence between other Cas13d orthologs from our new metagenomic mining (∼29.4% on average), we expect that our guide design model may generalize to other Cas13d effectors as well.

### DjCas13d exhibits high transcriptome-wide specificity

The context-dependent cellular toxicity mediated by many Cas13 enzymes is hypothesized to result from collateral cleavage of bystander transcripts (Buchman et al. 2020; Özcan et al. 2021; Ai et al. 2022; Shi et al. 2023). This is consistent with the observation that cellular viability and proliferation are more noticeably impacted when targeting more abundant transcripts – which would result in a larger number of activated Cas13 enzymes per cell and therefore more potential collateral RNA cleavage.

To investigate this hypothesis and compare the collateral and off-target effects between CasRx and DjCas13d, we performed RNA-seq two days after CasRx or DjCas13d-mediated knockdown of *CD81* (307 Transcripts Per Million (TPM)), *FTH1* (1219 TPM) and *ACTG1* (3728 TPM) in HEK293FT cells (**Figure 5A**). Our transcriptome-wide analysis revealed significantly more non-target transcripts affected by CasRx when targeting more highly expressed transcripts (*ACTG1*>*FTH1*>*CD81*), indicating greater levels of collateral or off-target effects (**Figure 5A**). In contrast, we observed minimal transcriptome-wide perturbation by DjCas13d apart from knockdown of the intended target transcript (**Figure 5A**).

**Figure 5:**
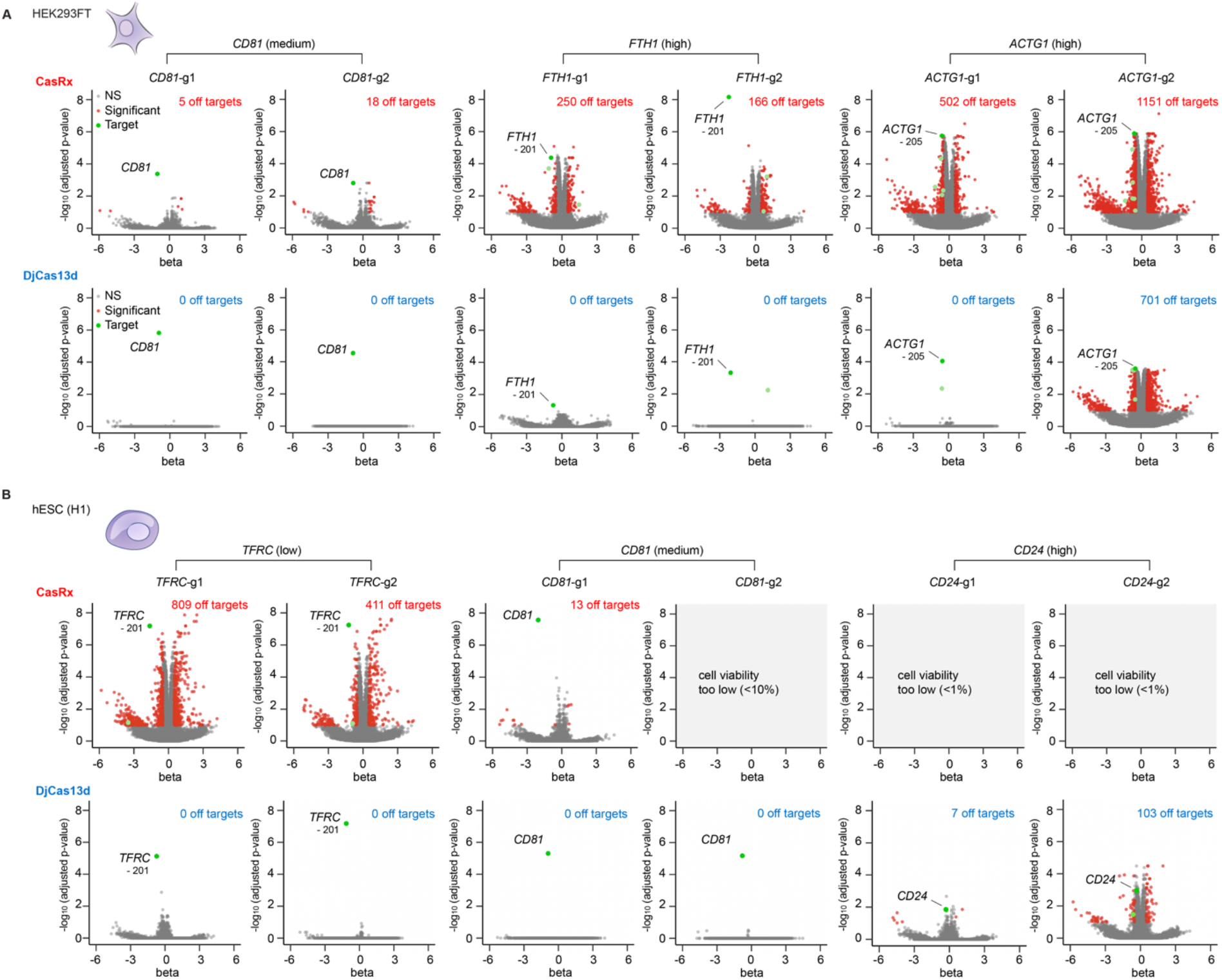
DjCas13d exhibits high transcriptome-wide specificity. **A.** Volcano plots of differential transcript levels between targeting guide conditions and non-targeting (NT) guide control for CasRx (top) and DjCas13d (bottom) in HEK293FT cells using two top-scoring guides for each target transcript (*CD81* (medium expression level)*, FTH1* (high expression level), and *ACTG1* (high expression level)). Red dots denote significantly affected transcripts with adjusted p value < 0.1 and beta value > |0.5|. Green dots denote target transcript isoforms, with darker green dots denoting the most abundant target transcript isoform, and lighter green dots denoting other significantly changed target transcript isoforms. N=3 biological replicates. **B.** Volcano plots of differential transcript levels between targeting guide conditions and non-targeting (NT) guide control for CasRx (top) and DjCas13d (bottom) in hESC (H1) cells with two top-scoring guides for each target transcript (*TFRC* (low expression), *CD81* (medium expression) and *CD24* (high expression)). Red dots denote significantly affected transcripts with adjusted p value < 0.1 and beta value > |0.5|. Green dots denote target transcript isoforms, with darker green dots denoting the most abundant target transcript isoform, and lighter green dots denoting other significantly changed target transcript isoforms. N=3 biological replicates.

Next, we extended our RNA-seq analysis to assess consequences of CasRx and DjCas13d in more sensitive hESC cells when targeting genes with high (*CD24*), medium (*CD81*), or low (*TFRC*) expression levels. CasRx-mediated knockdown of high and medium expressed genes resulted in rampant loss of cell viability, making transcriptome analysis impossible in many samples. Consistent with the high survival of sensitive cell types following DjCas13d treatment above, this toxicity was not observed for DjCas13d targeting the same transcripts. Similar to the HEK293FT RNA-seq above, we observed a significant reduction in off-target transcriptome perturbations when using DjCas13d (0 off-targets for most guides tested, with a modest 7 and 103 off-targets for the two guides targeting *CD24*) compared to CasRx (hundreds of off-targets even when targeting low- and medium-expression transcripts, and rampant cellular toxicity when targeting highly expressed transcripts) (**Figure 5B**).

Importantly, in order to rule out transcriptome-wide depletion that would be difficult to detect via differential RNA-seq, we used defined concentrations of exogenous RNA spike-ins to assess total RNA amount per cell. While CasRx showed a significant decrease in total RNA abundance across guides targeting *CD71*, DjCas13d did not display significant global RNA depletion with any guide/target tested, consistent with its low off-targets and low toxicity (**Figure S10A**). As an additional measure of transcriptome integrity, we visualized total RNA extracted from these samples and showed that while RNA integrity for DjCas13d was intact, CasRx targeting resulted in the appearance of a smaller molecular weight band between the 28S and 18S for all targeting guides (**Figure S10B**), which has also been noted by other groups (Shi et al., 2023).

To distinguish between guide-specific off-target effects and universal sequence-indiscriminate collateral effects in our CasRx datasets, we analyzed the overlap between up- and down-regulated transcripts among different guides, targets and cell types (**Figures S10C, D, E, F)**. We found a meaningful overlap between the significantly upregulated transcripts across different CasRx conditions, with enrichment of the unfolded protein response signaling pathway, suggesting that CasRx mediated non-target-specific collateral activity may stimulate generalized cellular stress responses.

### DjCas13d is a effective tool for gene knockdown in many sensitive cell types

Given the promise of DjCas13d as a high-fidelity and low-toxicity RNA targeting tool, we sought to apply DjCas13d to RNA targeting in sensitive biological processes and therapeutically-relevant cell types. Our demonstration of CasRx toxicity in hESC cells led us to assess DjCas13d knockdown in the context of hESC differentiation into neuronal progenitor cells (NPC), hematopoietic progenitor cells (HPC), and neurons. DjCas13d was delivered via an inducible Piggybac system at the stem cell stage and induced during differentiation. In NPCs, we targeted five transcripts including highly-expressed genes like *BSG* and *THY1*, and lower expressed transcripts such as *CD46* with one or two top-scoring guides per gene. We observed high cellular survival in all cases with no significant decrease relative to non-targeting conditions, and effective knockdown efficiencies in most cases, with a median of 63% (**Figure 6A**). In HPCs, we observed 46-69% knockdown of the target proteins CD81 and TRFC in DjCas13d-expressing cells with no detectable survival defect **(Figure 6B)**. In both of these cases, we confirmed that the expected markers of differentiation efficiency were not affected by DjCas13d targeting (SOX1 and PAX6 for NPC, CD43 for HPC) (**Figures 11A,B**). Finally, we differentiated hESCs to neurons using Neurogenin-2 (Ngn2) directed differentiation and assessed DjCas13d’s ability to knock down two proteins, CD81 and CD24, with 3 top-scoring guides each. We observed uniform knockdown of approximately 50% in all cases (measured at the protein level via FACS), coupled with high cell survival near 100% (median of 98%) (**Figure 6C**). Altogether, these data illustrate the broad applicability of DjCas13d across multiple target genes in sensitive cell types of high biological and therapeutic interest.

**Figure 6:**
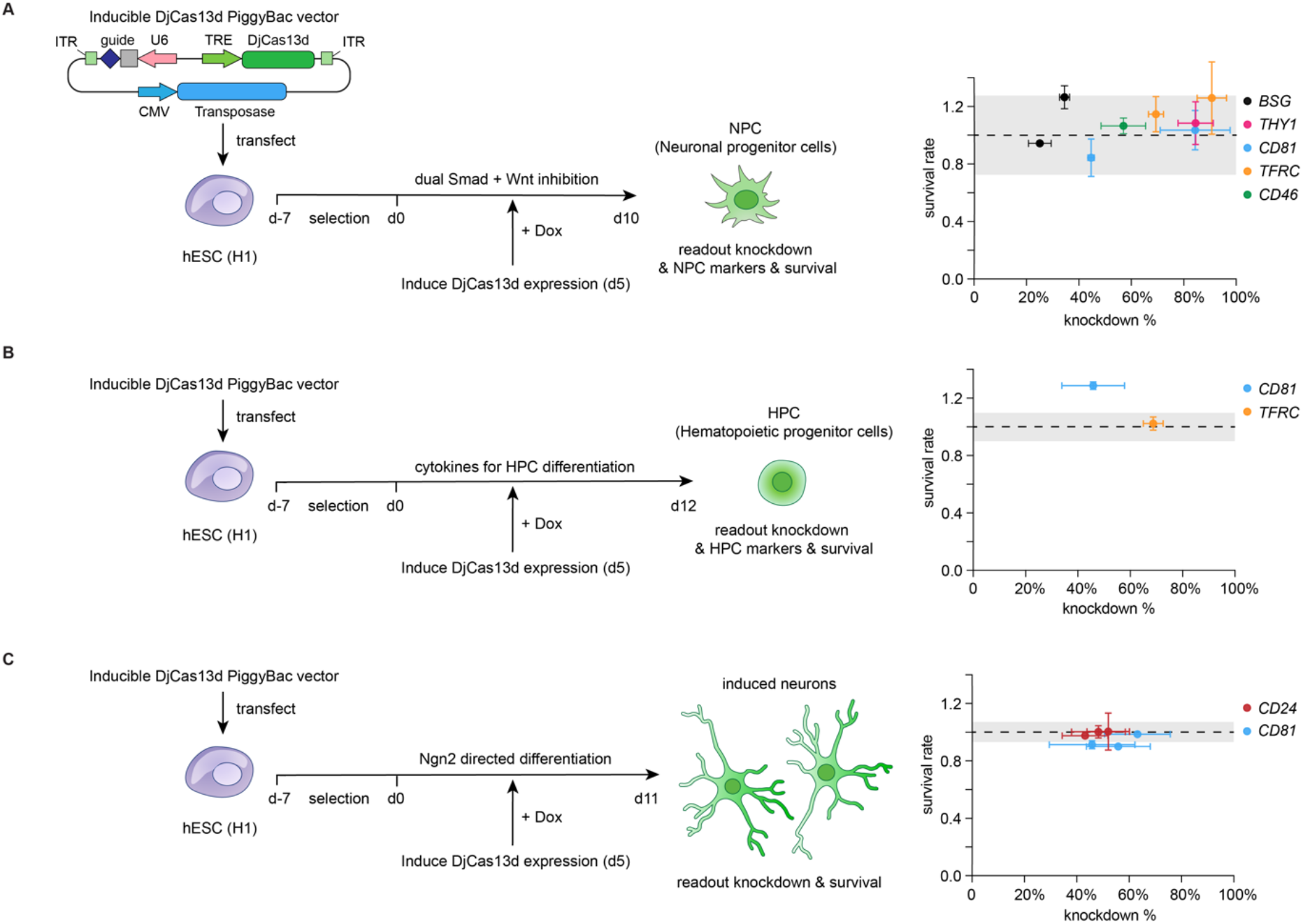
DjCas13d enables toxicity-free RNA perturbation in various sensitive cell types. DjCas13d-mediated RNA targeting in **A:** hESC-derived neuronal progenitor cells (NPCs); **B:** hESC-derived hematopoietic progenitor cells (HPCs); **C:** hESC-derived neurons. Left panel, schematic of the experimental workflow. Right panel, scatter plot of cellular viability (y axis) and knockdown efficiency (x axis) across five transcripts of different expression levels in NPCs and two transcripts of different expression levels in HPCs and neurons. Each dot on the scatter plot represents one guide’s survival rate and knockdown (mean ± SEM for n = 3 replicates). The dots are colored by the target transcript listed in the legend. The target transcripts are ranked by expression levels (high to low). The dots are colored by target transcripts. The black dashed line indicates a survival rate of 1.0 relative to the average of NT guides, and the shaded box indicates the SEM of the survival rate for NT guides.

To support easy use of both DjCas13d and CasRx for RNA targeting, we created a freely accessible portal to run our model for Cas13d guide prediction on all human and mouse transcripts and custom target sequences. This community resource is available at http://RNAtargeting.org.

## Discussion

In this study, we applied CasRx for large-scale screening across 127,000 guides against 55 target transcripts in human cells, a dataset that is >12 times larger than previous Cas13 guide design studies (Wessels et al. 2020; Cheng et al. 2023). Using this dataset, we developed a highly accurate, deep learning-based Cas13d guide efficiency model to nominate highly efficient guides for transcripts of interest. The model exhibits excellent performance across two screen modalities, nine cell types, and two diverse Cas13d orthologs, illustrating its generalizability for predicting highly effective guides across different contexts. The major factors contributing to our model’s generalizability include its primary reliance on the guide RNA spacer sequence - a cell type-independent feature - as well as the 9-fold cross-validation of the model on non-overlapping sets of transcripts, which alleviates overfitting to targeting hotspots specific to certain transcripts.

Previous attempts to predict CRISPR guide efficiency have primarily relied on manual selection of a limited set of guide sequence features combined with simpler machine learning models, such as elastic nets (Horlbeck et al., 2016), SVM (Doench et al., 2016), or random forest approaches (Wessels et al., 2020). More recently, deep learning models, which are able to learn complex, high-order patterns and features automatically from raw data, have been employed to predict guide efficiency for Cas9 activity (Chuai et al., 2018; Kim et al., 2019; Xue et al., 2019), Cpf1 (Kim et al., 2018), base editors (Arbab et al., 2020; Koblan et al., 2021), Cas13a (Metsky et al., 2022) and Cas13d (Cheng et al. 2023).

Here, we directly compared two deep learning models with linear and ensemble methods (elastic nets, random forest, and gradient-boosted trees) for guide efficiency prediction, finding that the deep learning model (CNN) outperformed the other approaches. This illustrates the power of deep learning models in sequence-based prediction tasks due to its automatic feature selection and ability to identify motifs or long-range interactions given a sufficiently large dataset (>100,000 guides). Furthermore, we show that our model significantly outperforms the current state-of-the-art models (Wessels et al. 2020; Cheng et al. 2023) (**Figures 3C, S8B**).

While deep learning models can extract important higher-order features automatically from raw inputs, the interpretation of feature contributions is challenging. Prior deep learning models for Cas9 (Chuai et al., 2018; Xue et al., 2019) and other sequence-based applications (Alipanahi et al., 2015; Kelley et al., 2016; Lanchantin et al., 2016) mainly employed neuron visualization methods to unveil important motifs. These approaches are able to successfully identify patterns recognized by individual filters, but can suffer from redundancy of the identified motifs. Recently developed interpretation methods such as Integrated Gradients, SHAP, and TF-MoDISco, can address these limitations and have begun to be applied to identify consolidated and non-redundant motifs for transcription factor binding (Avsec et al., 2021). In this report, we evaluated feature importance directly from the deep learning model using these new model interpretation approaches. This allowed us to discover a core region at guide spacer position 15-24 with a specific sequence composition predictive of high efficiency guides. Comprehensive motif analysis revealed a preference for GW_1-4_C_21_ or C_21_W_0-2_G motif. In contrast, analysis of base preference at individual positions and correlation-based evaluation of feature importance (Wessels et al., 2020) obscured this motif. This underscores the utility of the combination of deep learning models that are able to learn higher order sequence features along with advanced motif-discovery approaches for model interpretation such as TF-MoDISco used here – the first time, to our knowledge, that such an approach has been applied to CRISPR guide activity prediction models.

In addition to effective guide selection, cellular toxicity has emerged as a significant challenge for Cas13 applications, effects likely mediated by off-target and/or collateral RNA cleavage (Buchman et al. 2020; Özcan et al. 2021; Ai et al. 2022; Shi et al. 2023). Initial reports developing diverse Cas13 effectors for mammalian transcript knockdown demonstrated high specificity and lack of apparent cellular toxicity in HEK293FT cells, plants, and animal embryos (Abudayyeh et al., 2017; Cox et al., 2017; Konermann et al., 2018; Kushawah et al., 2020; Mahas et al., 2019). However, several recent studies have reported marked cellular toxicity of these effectors in other cell types or target contexts (Buchman et al., 2020; Özcan et al., 2021).

Two recent studies aiming to reconcile these reports concluded that collateral RNA cleavage by Cas13 enzymes is correlated with the expression level of the target transcript, and that the effect on cellular toxicity is dependent on the cell type (Ai et al. 2022; Shi et al. 2023), indicating that highly expressed transcripts and sensitive cell types are prone to Cas13-mediated collateral cleavage and toxicity. Our data comparing CasRx’s effect across cell types and endogenous target RNAs with varying expression levels supports this conclusion. We reasoned that more robust CasRx RNase activation upon higher target transcript levels would result in a greater amount of collateral RNA cleavage, which in turn could activate cellular stress pathways and lead to toxicity.

To advance Cas13 applications in sensitive cell types and therapeutic scenarios, our discovery of the DjCas13d ortholog promises to address current limitations of both CasRx (context-dependent cellular toxicity) and Cas7-11 (efficiency and size). DjCas13d exhibits minimal cellular toxicity even in challenging conditions, and achieves high efficiency and transcriptome-wide targeting specificity against highly expressed transcripts across various cell types. We further demonstrate efficient and high-viability endogenous RNA targeting with DjCas13d in hESC-derived neuronal progenitor cells (NPCs), hematopoietic progenitor cells (HPCs), and neurons. Therefore, DjCas13d is poised to overcome the limitations of previous tools. Future work characterizing mechanistic distinctions between CasRx and DjCas13d may reveal further protein engineering opportunities.

Taken together, DjCas13d paired with our state-of-the-art Cas13d guide design model provides a comprehensive solution for 3 key challenges in the RNA targeting toolbox by enabling high efficiency, cell viability, and specificity. We further envision that the deep learning model architecture, systematic feature engineering, and model interpretation approach outlined in this study will be broadly applicable to other sequence-based tasks, such as the prediction of guide RNA activities for newly discovered CRISPR enzymes, DNA/RNA modifications, and DNA/RNA-protein interactions.

## Data and Code Availability

The model is freely accessible at http://RNAtargeting.org. The RNAseq data is available at the NCBI Sequence Read Archive (SRA): PRJNA857683.

## Supporting information

Table S1. Oligos and primers

Table S2. Essential gene list

Table S4. Cas13d sequences

Table S3. Individual guide sequences

## Acknowledgments

We thank the Konermann laboratory and Hsu laboratory for support and advice; A. Pawluk for help with the manuscript; A. Kundaje and J. Zou for advice on deep learning models; J. Zou for the recommendation of the LinearFold package; W. Zhuk for building the initial deep learning model architecture; HK. Wayment-Steele for advice on RNA structure and energy prediction; A. Shrikumar for advice on the implementation and application of TF-MoDISco to our analysis; The Salk Institute NGS core and the Stanford Shared FACS facility for their support; and B. Hsu for helping build the CasRx guide design website. K.F. was supported by a UCSD Eureka! Scholarship for this work. S.K. is a Hanna Gray Fellow of the Howard Hughes Medical Institute, a Chan Zuckerberg Biohub Investigator, and an Arc Institute Core Investigator. P.D.H. is supported by the NIH (DP5 OD021369, R01 GM131073, R01 GM132465), DARPA, Emergent Ventures, the Shurl and Kay Curci Foundation, and the Rainwater Charitable Foundation and the Arc Institute.

## Author Contributions

S.K. and P.D.H. conceived this study and supervised the design and analysis of all experiments. J.W. and H.K. built the computational models, performed feature engineering, and implemented model interpretation. S.K. and J.W. analyzed the NGS data from the screens and calculated secondary features. J.W. performed the validation screen and individual guide testing in cancer cells. J.W. created the Cas13d guide efficiency prediction tool and performed model comparison. S.K. and P.D.H. computationally identified novel Cas13d orthologs. P.L. and E.W. cloned all Cas13d orthologs and tested in HEK293FT. J.W. tested the top Cas13d orthologs in stem cells. J.W., S.B., E.G., H.S., and E.K. cloned individual CasRx and DjCas13d guides. J.W., S.B., E.G., H.S., and E.K. performed individual guide testing in HEK293FT and stem cells. J.W. performed RNA-seq experiments and analyzed the data with C. V. D.. J.W., H.S., E.K., S.B., and E.G. performed RNA knockdown experiments in stem cell-differentiated neuronal progenitor cells, neurons, and hematopoietic progenitor cells. S.C. analyzed Cas13d ortholog sequences. M.D. performed computational mining of additional Cas13 sequences and built the Cas13 phylogenetic tree. P.L. S.K., and P.D.H. adapted CasRx for high-throughput screening. S.K., K.F., P.L., and P.D.H. performed the cell proliferation screen. J.W., S.K., and P.D.H. wrote the manuscript with input from all authors.

## Methods

### Plasmid design

For the CasRx expression vector, we designed a piggyBac-based all-in-one plasmid containing the CasRx effector, piggyBac transposase, and antibiotic selection cassette: PB_EF1a-CasRx-msfGFP-2A-Blast. The CasRx effector is fused to msfGFP at the C terminus and under the control of a constitutive EF1a promoter. A nuclear localization signal SV40 NLS was added to both the N and C terminus of CasRx-msfGFP. The antibiotic selection cassette, blasticidin S deaminase, is linked with CasRx-msfGFP via a P2A self-cleaving peptide.

For the CasRx guide cloning vector, we designed a lentiviral vector: hU6-(CasRx DR)-EF1a-Puro-WPRE. The CasRx DR is a 36-base direct repeat (CAAGTAAACCCCTACCAACTGGTCGGGGTTTGAAAC) for CasRx pre-gRNA (Konermann et al., 2018). The 30 nt guide spacer sequence is cloned into the vector through Gibson cloning using two BsmBI cleavage sites. For individual guide truncation and individual guide validation experiments, we designed a piggyBac-based all-in-one plasmid containing the CasRx effector, guide DR, piggyBac transposase, and antibiotic selection cassette: hU6-(CasRx DR)-TRE-CasRx-msfGFP-EF1a-rtTA-2A-Puro-CMV-transposase.

### Guide library design

For the survival screen, we selected 55 essential genes from the intersection of the essential hits in three previous survival screens performed in K562 cells (Hart et al., 2015; Horlbeck et al., 2016; Luo et al., 2008). We selected the major transcript isoform of these genes from the Refseq database and designed guides that tile these transcripts with single nucleotide resolution. A total of 127,071 targeting guides were generated for the 55 essential transcripts. In addition, we designed 14111 guides tiling 5 non-essential control transcripts (*CTCFL, SAGE1, TLX1, DTX2, OR2C3*). Along with 3563 non-targeting guides, we constructed a pooled library of 144745 guides.

For the validation screen on cell surface markers, 3218 guides were designed that tiled *CD58* transcripts (NM_001779.3, NM_001144822.2) and *CD81* transcripts (NM_004356.4, NM_001297649.2) with single nucleotide resolution. The targeting guides were pooled with 1186 non-targeting guides to create the final library.

### Guide library synthesis, cloning, and library amplification

For each guide spacer sequence in the guide library, we added a constant left overhang (“AACCCCTACCAACTGGTCGGGGTTTGAAAC”) and a right overhang (“TTTTTTTTGAATTCAAGCTTGGCGTAACTAGA”) to facilitate cloning. The resulting libraries were synthesized as oligo pools by Twist Biosciences, and then PCR amplified using the primer pair: Lib_F (“TCTTGTGGAAAGGACGAAACACCGCAAGTAAACCCCTACCAACTGGTCGGGGTTTG”) and Lib_R (“AGAGCTAGCCAGACGTGTGCTCTTCCGATCNNNNNNNNNTCTAGTTACGCCAAGCTTGA ATTC”) (**Table S1**). The PCR reaction was performed using NEBNext High Fidelity PCR Master Mix (NEB, catalog no. M0541L) for 20 cycles. The amplified library was gel-purified and cloned into the BsmBI digested guide cloning vector (hU6-(CasRx DR)-EF1a-Puro-WPRE) through Gibson assembly. The cloned guide library was then purified and concentrated by isopropanol precipitation.

For guide library amplification, the library plasmid was electroporated to Endura electrocompetent *E. coli* cells (Lucigen, catalog no. 60242-2) at 50–100 ng/ul. After electroporation, cells were recovered in LB medium for 1h, and then plated on LB agar plates with 100 ug/mL carbenicillin at 37°C for 12-14h. The colonies were then harvested at a coverage of > 500 colonies per guide. The amplified guide library plasmid was extracted using the Macherey-Nagel NucleoBond Xtra Maxi EF Kit (Macherey-Nagel, catalog no. 740424.10). To determine guide RNA representation, we PCR amplified the guide region using customized NGS primers containing Illumina adaptor sequences (**Table S1**). NextSeq sequencing was performed to determine guide RNA representation in the guide library. We verified that the library had >87% perfectly matching guides, <0.5% undetected guides, and a skew ratio (90th percentile:10th percentile read number) of less than 10.

### Lentivirus production

To produce lentivirus for the guide library, HEK293FT cells, purchased from Thermo Fisher (Cat # R70007) were grown in DMEM supplemented with 10% FBS (D10 media) at 37 °C with 5% CO2. Cells were passaged at a ratio of 1:2 using TrypLE (Gibco) and seeded 20–24 h before transfection at 1.8 × 10^7^ cells per T225 flask. For lentiviral plasmid transfection, the guide library plasmid was mixed with psPAX2 (Addgene, catalog no. 12260) and pMD2.G (Addgene, catalog no. 12259) in Opti-MEM, and transfected to HEK293FT using Lipofectamine 2000 (Thermo Fisher, catalog no. 11668027) and PLUS reagent (Thermo Fisher, catalog no. 11514015).

Medium was replaced 4 hours after transfection with fresh, prewarmed D10 medium. Two days after the start of lentiviral transfection, the supernatant from the HEK293FT cells was harvested and filtered using a 0.45um Stericup filter. The lentiviral titer was determined through spinfection on K562 cells prior to the screen.

### Cell culture and CasRx cell line generation

K562 cells were purchased from ATCC (CCL-243), and cultured in RPMI 1640 medium with GlutaMAX™ supplement (Thermo Fisher, catalog no.61870036), 10% FBS, and Penicillin-Streptomycin at 37 °C with 5% CO2. To generate a stable CasRx-expressing K562 cell line, we transfected K562 cells with the piggyBac-based all-in-one CasRx expression vector (PB_EF1a-CasRx-msfGFP-2A-Blast) using Lipofectamine 3000 Transfection Reagent (Thermo Fisher, catalog no. L3000001). Two days after transfection, we selected the cells with 10 μg/ml blasticidin S (Thermo Fisher, catalog no. A1113903). After selection for 1-2 weeks, we checked the percentage of CasRx-expressing cells using flow cytometry and confirmed that more than 95% of cells expressed CasRx-GFP.

### Survival screen

The guide library for the survival screen was lentivirally transduced at MOI=0.2 by spinfection into the stable CasRx-expressing K562 cell line. We ensured the guide library had a coverage of >1000 cells per guide. Two days after transduction, cells were selected with 1 μg/ml puromycin to ensure guide expression and further cultured for 14 days. Cells were harvested at day 14 (end of the screen), and the genomic DNA was extracted using Zymo Research Quick-gDNA MidiPrep (Zymo Research, cat. no. D4075). The guide region was PCR amplified using customized NGS primers containing Illumina adaptor sequences. The resulting PCR products were gel purified and quantified with Nanodrop and Qubit dsDNA HS Assay Kit (Thermo Fisher Scientific, cat. no. Q32851). Pooled guide libraries were sequenced on Illumina NextSeq, with 80 cycles of read 1 (forward) and 8 cycles of index 1. Three biological replicates were performed for the survival screen.

### Validation screen on *CD58* and *CD81*

The guide library for the validation screen on *CD81* and *CD58* was lentivirally transduced at MOI=0.1 by spinfection into the stable CasRx-expressing K562 cell line. 45 million cells per biological replicate were transduced to ensure coverage of >1000 cells per guide. After spinfection, cells were selected with 1 μg/ml puromycin and further cultured for 10 days. At the end of the screen, cells were divided into two pools and stained with *CD58* antibody (BD Biosciences, catalog no. 564363) or *CD81* antibody (BD Biosciences, catalog no. 561958) and analyzed using FACSAria II (**Figure S11**). Following calibration with unstained controls, each cell pool was sorted into four bins based on target gene expression level indicated by antibody-conjugated fluorescence intensity. Specifically, cells were first gated by forward and side scatter to select for live, single cells. Next, cells were gated on GFP to select for CasRx-expressing cells. This final population was sorted into four bins based on the intensity of *CD58* or *CD81* antibody-conjugated fluorescence intensity. As high efficiency guides were defined as the top 20% for each gene, we set the bin with the lowest target gene expression (bin 1) at 7-8%, which is equal to the fraction of the target gene’s high efficiency guide number in the whole library: 1600*0.2/4401. The rest of the population was equally divided into three bins of the same size (∼30%). The genomic DNA for cells in each bin was extracted and sequenced as in the survival screening. Four biological replicates were performed for the validation screen.

### Data preprocessing and definition of high efficiency guides

For each guide RNA we calculated the fraction in the day 14 guide pool and the input library pool. Guide efficiency was evaluated by the ratio of guide percentage in the day 14 pool to the input pool (**Table S3**). Guides targeting each transcript were ranked based on their average ratio across the three replicates, and we defined high efficiency guides for each transcript, taking into account three parameters: 1) the top 20% guides per transcript; 2) no essential off-targets predicted by BLAST (see the ‘Off-target filtering’ section below and **Figure S1D** for details); and 3) with a d14/input ratio lower than 0.75 (the ratio at 5th percentile of the guides for targeting control genes). Guides targeting the transcript *RPS19BP1* were excluded because they clustered with non-essential controls (most guides were not effectively depleted in the screen).

For the validation screen, we first filtered guides with less than 200 counts in all *CD58* bins and *CD81* bins. Less than 1.5% of guides were removed by this filter. We then calculated each guide’s distribution across the 4 bins and used the ratio of guide percentage in bin 1 (greatest knockdown) to the sum of its percentage in bins 1 and 4 (least knockdown) for evaluation of guide efficiency. We then ranked guides within each gene based on their average ratio of the four replicates, and we defined the top 20% guides for each gene as high efficiency guides.

### Off-target filtering

We performed BLAST to identify potential off target matches for our guides. As the first 24 nucleotides from the 5’ end of the CasRx guides were shown to be most indicative of guide targeting ability (**Figure S2C**), we took the first 24 nucleotides of each guide as BLAST input. BLAST was performed using a generous E value of 1 (e=1) against the Gencode V33 database. BLAST results were parsed and off target genes were identified as those with up to three mismatches to the guide input. To check the essentiality of the off-target genes, we made an essential gene list by combining the essential gene hits from the three previous survival screens in K562 cells and we compared the off-target genes with the essential gene list. Guides with predicted off-targets in essential genes were filtered, as we reasoned they may interfere with the interpretation of our survival screen readout. For our survival screening, 6790 guides were filtered and 120281 guides remained for further analysis (**Figure S1D**, Table S4). For the validation screen, the filtered dataset is provided in Table S5.

### Analysis of the positional distribution of high efficiency guides

For each transcript, we calculated the number of high efficiency guides overlapping each position on the transcript, and plotted the results using a heatmap. We further summarized the distribution of high efficiency guides across all transcripts and positions with a histogram. In theory, a particular nucleotide position would have at most 30 guides covering it, so the number of efficient guides ranges from 0 to 30 for each position. We compared the results with a randomly sampled distribution, which is simulated using 100 random samplings of 20% of the guides in the library. In theory, the randomly sampled distribution would show a peak at 6 (30*20%), which agrees with our simulation results.

### Data splits

For model hyperparameter tuning and evaluation, we split our 54 essential transcripts into 9 folds, each containing a unique and non-overlapping set of 6 test transcripts. The 54 transcripts were distributed evenly across the 9 folds according to their high efficiency guide percent to make the 9-fold split balanced. Using the predefined transcript splits, we performed 9-fold cross-validation to tune model hyperparameters and compare prediction accuracy between models.

### Feature calculation and model inputs

For the sequence input, each 30 nt guide spacer was one-hot encoded into four binary vectors of length 30 to represent the nucleotide identity at each position.

To predict guide unfolding energy, we used LinearFold, a linear-time RNA secondary structure prediction algorithm (Huang et al., 2019) on the full-length guide sequence (36nt DR +30nt spacer). We started with the default parameters and the CONTRAfold v2.0 model (Do et al., 2006; Lorenz et al., 2011; Wayment-Steele et al., 2020) provided by the LinearFold software at https://github.com/LinearFold/LinearFold. We subtracted the predicted MFE (minimum free energy) with the baseline energy (MFE of the unstructured guide with the 30 nt spacer unfolded) to calculate guide unfolding energy. We also tested the Vienna RNAfold model in LinearFold as a comparison. To determine whether using the ensemble guide unfolding energy instead of MFE could improve model prediction, we further tested three RNA structure prediction algorithms (Contrafold2, Eternafold, Vienna) wrapped by Arnie (https://github.com/DasLab/arnie) to calculate the ensemble guide unfolding energy with the partition function (Do et al., 2006; Lorenz et al., 2011; Wayment-Steele et al., 2020). For the Vienna package, we tested different temperature(T) settings: 37°C, 60 °C, and 70 °C. In our final model, we used the guide unfolding energy calculated by LinearFold’s default CONTRAfold v2.0 model as it improved model prediction accuracy to the greatest extent.

To calculate target unfolding energy, we first used LinearFold’s CONTRAfold v2.0 model to predict MFE of the native local target region using the local target sequence. We then predicted MFE of the guide unwound local target region by supplying the algorithm with the constraint that the 30 nt guide-binding site is unpaired. (This can be achieved by feeding in an additional constraint structure with the guide-binding site annotated with “.”). We then subtracted the former MFE (MFE of the native target region) by the latter (MFE of the guide unwound target region) to estimate local target unfolding energy. The local target region was defined as the 30 nt guide-binding site with 15 nt flanking sequence on both sides. Flanking sequences of different lengths were compared, and the length 15 was chosen for the final model as it improved model prediction accuracy to the greatest extent.

To calculate the percentage of isoforms targeted by each guide, we obtained all transcript isoforms for each gene from the Refseq database and evaluated the percentage of isoforms matched for each 30nt guide target (using perfect matches).

To calculate the three position flags, we obtained Refseq’s annotations of the 5′ UTR, CDS, or 3′ UTR region for our target transcripts. Guides that target the 5′ UTR, CDS, or 3′ UTR region have a flag value of 1 for that correspondent feature, and 0 for the other two flag features. To calculate the three position floats (5′ UTR position,CDS position,3′ UTR position), we calculated the relative position of the guide target site in the 5′ UTR, CDS, or 3′ UTR region. Guides located out of the region have a flag value of 0 for the correspondent feature.

### Model architecture

#### Sequence-only models

For linear models and ensemble models, the one-hot encoded guide sequence was flattened and converted to 30*4= 120 flag features. The features are then fed into the models to generate the output. For the CNN model, the one-hot encoded guide was treated as a 4-channel image, and a few 1D convolutional layers were applied to generate a feature map, which was flattened and passed to a dense layer to generate the final output. For the biLSTM model, the guide sequence was treated as a sentence with four characters, and two LSTMs, each processing the input sequence in one direction (forward or backward), were applied to generate sequence representations. The resulting vectors were merged, flattened, and passed to a dense layer to generate the final output.

#### Full model with secondary features

For the CNN model with secondary features, the one-hot encoded guide was passed to a few convolutional layers as in the sequence-only model. The output from the CNN layers was flattened and concatenated with the normalized secondary features. The concatenated feature vector was sequentially passed to a dense layer, a recurrent dense layer and a final dense layer of 1 unit to generate the output. All dense layers use leaky ReLU as the activation function. The CNN layer kernel size, unit number, layer number and the dense layer unit number were defined after hyperparameter tuning.

For the Gradient-boosted classification tree, the one-hot encoded guide sequence was flattened and converted to 30*4= 120 flag features. The sequence features are concatenated with the normalized secondary features, and then fed into the model to generate output.

### Model training, hyperparameter tuning and evaluation

All models were trained to solve a binary classification task – predicting probabilities of high efficiency guides. The linear models and ensemble models were trained in scikit-learn 0.24 and the deep learning models (LSTM and CNN) were trained in TensorFlow 2.3.1. For the deep learning models, we used binary cross-entropy as the loss function and applied the Adam optimizer for model training. Early stopping was used to prevent model overfitting. For all models, the prediction accuracy is evaluated by AUROC (Area Under the Receiver Operating Characteristic curve) and AUPRC (The Area Under Precision-Recall Curve).

To tune hyperparameters and evaluate model performance, we used 9-fold cross-validation over the hyperparameter space. For linear models and ensemble models, we used the “GridSearchCV” function in scikit-learn to perform a grid search over the hyperparameter set. For deep learning models, we used the Hyperband tuner in TensorFlow to select top models quickly by filtering out poor models during training.

The hyperparameter sets for all models are listed below:

- logistic regression with L1 regularization: regularization strength in logarithmic intervals from 10^−5^ to 10^5^
- logistic regression with L2 regularization: regularization strength in logarithmic intervals from 10^−5^ to 10^5^
- logistic regression with elastic net regularization: regularization strength in logarithmic in intervals from 10^−4^ to 10^4^, L1 ratio equally spaced from 0.1 to 1.
- Gradient-boosted classification trees: number of trees – [100,200,400,800,1000,1200,1500,1800,2000], maximum depth of a tree – [2,4,8], number of features to consider when looking for the best split – [all, sqrt(n_features), log2(n_features)].
- Random forest (RF): number of trees – [100,200,400,800,1000,1200,1500,1800,2000], number of features to consider when looking for the best split – [all, sqrt(n_features), log2(n_features)].
- Long short-term memory recurrent neural network (LSTM): LSTM units – [16, 32,64,128], dense layer units – [8, 16, 32], number of recurrent dense layes – [0,1,2,3], dropout rate – [0.0, 0.1, 0.25]
- Convolutional neural network (CNN): CNN layer kernel size – [3,4,5], CNN unit – [8,16,32,64], number of CNN layers – [3,4,5], number of dense layer units – [8,16,32,64] number of recurrent dense layers – [0,1,2,3]

For all models, we chose the hyperparameter set with the highest average AUROC across all test sets among the 9-fold splits, and evaluated the final model performance using both the average AUROC and average AUPRC across test sets.

### Secondary feature selection

For the CNN model, we added each secondary feature individually to guide sequence features and calculated the change in model performance. We selected features that successfully improved model performance, and added these features sequentially upon guide sequence features to check feature redundancy. We also tried removing individual features from the final model to confirm the necessity of the features.

For the Gradient-boosted tree, besides the above methods, we also used Boruta, an all-relevant feature selection method that aims to find all features useful for prediction (Kursa et al., 2010). We implemented it using BorutaPy, the Python implementation of Boruta (https://github.com/scikit-learn-contrib/boruta_py) on our Gradient-boosted tree.

### Model interpretation and feature contributions

For the CNN model, we applied “Integrated Gradients” (IG) to investigate feature contributions in the model. “Integrated Gradients” is an attribution method that evaluates feature importance by integrating the gradient of output to input features along the straightline path from the baseline input to the actual input value (Sundararajan et al., 2017). Due to the non-linearity of the deep learning model, we applied “Integrated Gradients” to the best-performing individual CNN model on CD genes rather than the ensemble model. To compute integrated gradients, we first set all-zero baselines for the sequence input, position flags and position floats, and used average baselines for other features. Next, we generated a linear interpolation between the baselines and the inputs using 50 steps. We then computed gradients using the “tf.GradientTape” function in TensorFlow for the interpolated points, and approximated the gradients integral with the trapezoidal rule. To evaluate the relative importance of each position on the guide, we averaged the absolute integrated gradient values at each position across all test sequences. To evaluate the contribution of each nucleotide at each position, we averaged the integrated gradients for that nucleotide across all test sequences.

For the Gradient-boosted tree, we applied SHAP (SHapley Additive exPlanations) to investigate feature contributions in the model. SHAP is a game theoretic approach that estimates how each feature contributes to the model output by providing the SHAP value for each input feature (Lundberg et al., 2020). We implemented the SHAP package from https://github.com/slundberg/shap, and applied it to our Gradient-boosted tree. To evaluate the relative importance of each position on the guide, we averaged the SHAP values at each position across test sequences. To evaluate the contribution of each nucleotide at each position, we averaged the SHAP values for that nucleotide across test sequences.

### Cas13a guide sequence contribution to guide efficiency

We analyzed three Cas13a guide efficiency datasets: 1) the Luciferase knockdown dataset containing 186 LwaCas13a guides for *Gaussia* luciferase (Gluc) and 93 guides for *Cypridina* Luciferase (Cluc) (Abudayyeh et al., 2017); 2) the endogenous gene knockdown dataset containing 93 LwaCas13a guides for each of *KRAS*, *PPIB* and *MALAT1* (Abudayyeh et al., 2017); and 3) the ADAPT dataset containing 85 perfect match LwaCas13a guides for virus detection (Metsky et al., 2022). We calculated the Pearson correlation between each nucleotide at each position with guide efficiency to evaluate the sequence contribution.

### Motif discovery

For motif discovery, we used TF-MoDISco (Transcription Factor Motif Discovery from Importance Scores), an algorithm that discovers motifs by clustering important regions in sequences using per-base importance scores (Shrikumar et al., 2018). We implemented TF-MoDISco from https://github.com/kundajelab/tfmodisco using the integrated gradients of all high efficiency guides in our training data as input. We ran TF-MoDISco with a sliding window size of 7 and a flank length of 2. For final motif processing, we trimmed the clustered motifs to a window size of 6, added an initial flank length of 2 and a final flank length of 3 to get the final motifs. The top 5 active motifs are picked and aligned to the 30 nt spacer according to the mode position of sequences in each motif.

### Nmer analysis

To identify enriched or depleted positional nmers, we divided our survival screen data to 9 folds as in the model training workflow and calculated the ratio of all possible positional nmers’ percentage in high efficiency guides to non-high efficiency guides in the training set and test set, respectively, for each fold. We identified enriched (or depleted) nmers based on their ratio in the training set with a predefined ratio cut-off. We selected the nmers identified as enriched (or depleted) across all folds, and ranked them by their average percent in high efficiency guides in the test sets across all folds. The initial ratio cut-off is set as 2 for enriched nmers and 0.5 for depleted nmers. The cut-off is adjusted during the nmer identification process so that the percent of guides with enriched nmers are ∼20% and the percent of guides with depleted nmers are ∼40%. We mainly focused on 3-mers and 4-mers in this paper.

### Final model and model testing on the validation screens

We chose the CNN model as our final model after hyperparameter tuning and model comparison. We re-trained the model using all of the survival screen data. To prevent overfitting, we split out a validation set during model training as in the previous 9-fold cross-validation split. We built 9 individual models using different validation sets from the 9-fold split of essential transcripts, and we compared their performance on the two cell surface markers, *CD58* and *CD81*. We further built an ensemble model that averaged the prediction of all the individual models. We found that the ensemble model outperformed all individual models on the two CD genes, so we set the ensemble CNN model as our final model. As a comparison, we also retrained the best non-deep learning model, the Gradient-boosted tree (GBT), using all of the survival screen data. We tested the model on the two CD genes and evaluated model performance using AUROC and AUPRC.

### Model comparison with Wessels et al. model and DeepCas13

We tested the performance of the Random forest model from Wessels et al. on our CD genes and essential genes using the web server https://cas13design.nygenome.org (Wessels et al. 2020; Guo et al. 2021). We evaluated the model performance using AUROC, AUPRC, Spearman’s correlation coefficient, r_s_ and true positive ratios at 0.8 and 0.9 model score cutoffs. As the Random forest model is designed for 23 nt long guides, we extended the guides from their model output to 30 nt (extends toward the 3′ end) to be in accordance with our screen data. For comparison, we retrieved the CasRx guide tiling screen dataset on three genes, *CD46*, *CD55*, and *CD71*, from Wessels et al. and tested our model’s performance. We adjusted the guide length to 23 nt in our model to be in accordance with their screen data, and we set the top 20% guides for each gene as “high efficiency guides”. The model performance was also evaluated by AUROC, AUPRC, Spearman’s correlation coefficient, r_s_ and true positive ratios at 0.8 and 0.9 model score cutoffs.

We tested the performance of DeepCas13 <u>(Cheng et al. 2023)</u> on our CD genes using the web server http://deepcas13.weililab.org. We evaluated the model performance using AUROC, AUPRC, Spearman’s correlation coefficient, r_s_ and true positive ratios at 0.8 and 0.9 model score cutoffs.

### Cas13d guide efficiency prediction tool and website

A website-based Cas13d guide efficiency prediction tool was developed using our CNN model for Cas13d guide design across model organism transcriptomes and custom RNA sequences. For model organism Cas13d guide design, we precomputed the Cas13d guide efficiency for all coding and non-coding genes of each model organism. Briefly, reference transcriptome sequences and annotations were obtained from the UCSC Table Browser (Karolchik et al., 2004) with the NCBI RefSeq track. All possible 30 nt Cas13d guide spacers were extracted from the transcriptome sequences with single nucleotide resolution. Secondary features were calculated for each guide as described in the ‘**Feature calculation and model inputs’** section above. The final CNN model was applied to all guides for prediction of their efficiency, and the guides were ranked within each gene based on the model prediction scores.

For custom sequence guide design, all possible 30 nt Cas13d guide spacers are extracted from the input custom RNA sequences with single nucleotide resolution. Guide unfolding energy and target unfolding energy are calculated as described in the ‘**Feature calculation and model inputs’** section above. A CNN model that uses guide sequence, guide unfolding energy and target unfolding energy as inputs, trained on the survival screen dataset, is applied to the custom sequence guides for prediction of their efficiency. Guides are ranked based on the model prediction scores.

The Cas13d guide efficiency prediction tool is freely available on a public, user-friendly website: https://www.RNAtargeting.org.

### Computational identification of novel Cas13d orthologs through metagenomic database mining

We applied our previously described pipeline for novel CRISPR effector discovery (Konermann et al., 2018) to incompletely assembled metagenomic contigs in addition to whole genome, chromosome, and scaffold-level prokaryotic and metagenomic sample assemblies from the NCBI Genome database (https://www.ncbi.nlm.nih.gov/), the Gigadb repository (http://gigadb.org/), as well as the JGI Genome portal (https://genome.jgi.doe.gov/portal/).

Putative effectors encoded near identified CRISPR arrays (<kb distance) were assigned to previously identified Cas13 families via tBLASTn analysis, where a bit score of at least 60 to any prior Cas13 subfamily member was required for cluster assignment. As a second round of discovery independent of CRISPR array identification, tBLASTn was performed on all original and predicted Cas13d effectors from the first round against all public metagenome whole genome shotgun sequences without predicted open reading frames (ORFs) from all three sources listed above. New full-length homologs and homologous fragments were aligned using Clustal Omega and clustered using PhyML 3.2 (Guindon et al., 2010). All the Cas13d ortholog sequences are provided in **Table S7**.

### Construction of Cas13 phylogenetic tree

A custom sequence database of bacterial isolate and metagenomic sequences was constructed by aggregating publicly available sequence database, including NCBI, UHGG (Almeida et al., 2021), JGI IMG (I.-M. A. Chen et al., 2021), the Gut Phage Database (Camarillo-Guerrero et al., 2021), the Human Gastrointestinal Bacteria Genome Collection (Forster et al., 2019), MGnify (Mitchell et al., 2020), Youngblut et al animal gut metagenomes (Youngblut et al., 2020), MGRAST (Meyer et al., 2008), and Tara Oceans samples (Sunagawa et al., 2015). Cas13 sequences from other Cas13 families were identified by searching representative members of each clade (Cas13a/b/bt/c/x/y) against a collection of protein representatives (clustered at 30% identity) derived from the custom sequence database using hmmsearch from the hmmer package (*HMMER*, n.d.). Selected Cas13a, Cas13b, Cas13c, Cas13d representatives were LbuCas13a, BzoCas13b, AspCas13c, and CasRx respectively. The Cas13bt representative was collected from (Kannan et al., 2022), and the Cas13X and Cas13Y representatives were collected from (Xu et al., 2021). All hits that met E < 1e-6 and were 75%-125% the length of the representative sequence were retained. Sequences were assigned to the best matching representative. Sequences were then clustered at the 50% identity level along 80% of both sequences using the mmseqs package (Steinegger & Söding, 2017). Sequences were then aligned using the MAFFT algorithm mafft-linsi (Katoh et al., 2002). PhyML was used to generate phylogenetic trees with default parameters (Guindon et al., 2010). Trees were visualized using the ggtree package in R (Yu, 2020).

### Cloning of Cas13d orthologs and Cas7-11

For initial testing and efficiency screening, human codon optimized Cas13d sequences, flanked by two nuclear localization or export sequences, were cloned into a backbone derived from pXR001: EF1a-CasRx-2A-EGFP (Addgene #109049) to replace the CasRx coding sequence. Guide sequences targeting mCherry or *CD81* were cloned into a backbone derived from pXR003: CasRx gRNA cloning backbone (Addgene #109053) with 5′ full-length direct repeat (DR) sequences for each Cas13d ortholog. For testing the seven high efficiency Cas13d orthologs in stem cells, the Cas13d coding sequences and respective mature DR guide scaffold sites were cloned into the inducible piggyBac-based all-in-one plasmid containing the Cas13d effector, guide DR, piggyBac transposase, and antibiotic selection cassette: hU6-DR-TRE-Cas13d-T2A-msfGFP-EF1a-rtTA-T2A-Puro-CMV-transposase. Human codon optimized DisCas7-11 protein sequence and the mature DR guide scaffold with golden gate sites were PCR amplified from Addgene plasmids # 172507 and #172508, a gift from Omar Abudayyeh & Jonathan Gootenberg, and cloned to the constitutive piggyBac-based all-in-one backbone plasmid as mentioned before. Guide spacers were position matched to CasRx and DjCas13d’s guide spacers and were cloned into the backbone plasmid using Golden Gate cloning. All individual guide sequences are provided in **Table S6**.

### Cell culture for individual guide testing

HEK293FT cells were purchased from Thermo Fisher (Cat # R70007) and grown in DMEM supplemented with 10% FBS (D10 media) at 37 °C with 5% CO2. Cells were passaged at a ratio of 1:2 using TrypLE (Gibco). Hela and A375 cells were gifts from the Howard Chang lab and Scott Dixon lab, respectively. They were both cultured in DMEM supplemented with 10% FBS (D10 media) at 37 °C with 5% CO2. Cells were passaged at a ratio of 1:2 using TrypLE (Gibco). U2OS cells were a gift from the Chang lab and grown in McCoy’s 5A (modified) Medium (Thermo Fisher, catalog no. 11668027) supplemented with 10% FBS at 37 °C with 5% CO2. Cells were passaged at a ratio of 1:2 using TrypLE (Gibco). Stem cell line H1 were purchased from WiCell (Cat # WA01). Cells were maintained in mTeSR™ Plus media (Catalog # 100-0276, STEMCELL Technologies) on Matrigel-coated 6-well plate and passaged 1:12 with ReLeSR™ (Catalog # 05872, STEMCELL Technologies) every four days.

### Transfection of human cell lines

For initial testing and efficiency screening of Cas13d orthologs, HEK293FT cells were plated at 20,000 cells per well in a 96-well plate, then transfected at >80% confluence with 192 ng Cas13d-2A-EGFP plasmid, 192 ng of crRNA expression plasmid, and 12 ng of mCherry expression plasmid using Lipofectamine 2000. Cells were harvested 48 hours after transfection for flow cytometry analysis of mCherry expression. For CD81 knockdown experiments, HEK293FT cells were transfected with 200 ng Cas13d-2A-EGFP plasmid and 200 ng guide RNA expression plasmid using Lipofectamine 2000. Cells were harvested 48 hours after transfection for staining and flow cytometry analysis of CD81 expression.

For experiments comparing CasRx, DjCas13d, and Cas7-11 in HEK293FT cells, cells were plated at 16,000 cells per well in a 96-well plate and transfected at > 80% confluence with 100 ng of all-in-one PiggyBac plasmids containing CasRx, DjCas13d, or Cas7-11 using Lipofectamine 2000 (Life Technologies). Cells were selected with 1 μg/ml puromycin 24h after transfection. 24 hours after selection, cells were harvested for RNA extraction and downstream processing.

For individual guide testing in Hela cells, low passage cells were plated at a density of 15,000 cells per well in a 96-well plate and transfected at > 80% confluence with all-in-one PiggyBac plasmids containing CasRx or DjCas13d using FuGENE® HD Transfection Reagent (E2311, Promega) according to the manufacturer’s protocol. Cells were selected with 1 μg/ml puromycin and induced with Doxycycline (D3072, Sigma) for CasRx or DjCas13d expression 48h after transfection. Flow analysis was performed seven days after induction.

For individual guide testing in U2OS cells, low passage cells were plated at a density of 15,000 cells per well in a 96-well plate and transfected at > 80% confluence with all-in-one PiggyBac plasmids containing CasRx or DjCas13d using ViaFect™ Transfection Reagent (E4981, Promega) according to the manufacturer’s protocol. Cells were selected with 0.75 μg/ml puromycin and induced with Doxycycline (D3072, Sigma) for CasRx or DjCas13d expression 48h after transfection. Flow analysis was performed seven days after induction.

For individual guide testing in A375 cells, low passage cells were plated at a density of 25,000 cells per well in a 96-well plate and transfected at > 80% confluence with all-in-one PiggyBac plasmids containing CasRx using TransIT-X2 (MIR 6003, Mirus) according to the manufacturer’s protocol. Cells were selected with 0.5 μg/ml puromycin and induced with Doxycycline (D3072, Sigma) for CasRx expression 48h after transfection. Flow analysis was performed seven days after induction.

For enzyme comparison and individual guide testing in H1 cells, low passage cells were passaged with Accutase (Innovative Cell Technologies) and plated into a Matrigel-coated 96-well plate with mTESR media containing ROCK inhibitor Y-27632 (10 uM, Abcam) at 30,000 cells per well one day before transfection. On day 1, cells were transfected at > 80% confluence with all-in-one PiggyBac plasmids containing different Cas13d orthologs using FuGENE® HD Transfection Reagent (E2311, Promega) according to the manufacturer’s protocol. Cells were selected with 0.5 μg/ml puromycin 48h after transfection. 5-7 days after selection, Cas13d expression was induced with Doxycycline (D3072, Sigma). Flow cytometry analysis was performed three days after induction.

For RNAseq experiments in H1 cells, low passage cells were passaged with Accutase (Innovative Cell Technologies) and plated into Cultrex (R&D Systems 343400502)-coated 96-well plates with mTESR media containing ROCK inhibitor Y-27632 (10 uM, Abcam) at 25,000 cells per well one day before transfection. On day 1, cells were transfected at > 80% confluence with all-in-one PiggyBac plasmids containing different Cas13d orthologs using FuGENE® HD Transfection Reagent (E2311, Promega) according to the manufacturer’s protocol. Cells were split and selected with 0.75 μg/ml puromycin 24h after transfection. Puromycin concentration was increased to 1ug/ml the next day. 72h after transfection, cells were harvested for RNA extraction and downstream processing.

### Staining and flow cytometry

For cell surface protein staining, cells were harvested and dissociated with TrypLE, followed by two washes in cold FACS buffer (DPBS + 2 mM EDTA + 0.02% BSA), and then blocked with Human TruStain FcX (Biolegend) for 10 minutes. Cells were then stained with target antibodies for 1 hour at 4°C in the dark, followed by two washes using the FACS buffer, and then analyzed by flow cytometry.

For intracellular staining, cells were dissociated with Accutase and resuspended in DMEM/F12 with GlutaMAX (ThermoFisher, Cat #10565018) with 20% trypsin inhibitor. Cells were then fixed with Cytofix/Cytoperm solution (BD) at 4°C for 20 minutes, followed by washes with Perm/Wash solution (BD). Cells were then stained with target antibodies for 45 minutes at 4°C in the dark, followed by two washes with the FACS buffer, and then analyzed by flow cytometry.

### RT-qPCR

Cells were lysed with BME-supplemented RLT buffer and total RNA was extracted with the RNeasy Plus 96 Kit (Cat #74192, QIAGEN). The extracted RNA was then reverse transcribed using RevertAid RT Kit (Thermo Fisher, Cat # K1691) with random hexamer primers at 25°C for 5 min, 42°C for 60 min, and 70°C for 5 min. qPCR was then performed using Taqman Fast Advanced Master Mix (Thermo Fisher, Cat # 4444965) and Taqman probes for GAPDH control (Thermo Fisher, Cat # 4326317E) and target genes (IDT, custom gene expression assays).

Custom Taqman probe and primer sets were designed to amplify target regions spanning the guide target sites. qPCR was performed in 384-well plates using the LightCycler 480 Instrument II (Roche). Target gene expression change was calculated relative to non-targeting controls using the ddCt method.

### Cell viability assays

For cell viability assays in HEK293FT, cells were plated at 9,000 cells per well in a 96-well plate the day before transfection. Cells were transfected with 100 ng of all-in-one PiggyBac plasmid containing constitutive CasRx, DjCas13d, or Cas7-11 using Lipofectamine 2000 (Life Technologies). 72 hours after transfection, cell viability was measured using WST-1 reagent (5015944001, Sigma) with an incubation time of 2 hours and measurement of absorbance at 440nm. Cell viability of targeting guide groups for each effector was compared relative to the corresponding non-targeting guide group. Three biological replicates were performed.

To measure cell viability in stem cells, Hela, U2OS and A375 cells, cells were transfected with the inducible all-in-one PiggyBac plasmids containing inducible CasRx, DjCas13d, or other Cas13d orthologs. After selection for plasmid integration with 1 μg/ml puromycin for 5-7 days, cells were induced for effector (CasRx, DjCas13d or other Cas13d orthologs) expression using Doxycycline (D3072, Sigma). 3-5 days after induction, flow analysis was performed to quantify the percent of cells expressing the effector in each experimental group using the GFP reporter. The GFP+ percentage of cells with targeting guide groups for each effector was normalized to that of the corresponding non-targeting guide group for evaluation of cell viability upon target RNA knockdown. Three biological replicates were performed.

To measure cell viability in stem cell derived NPCs, HPCs, or neurons, we transfected stem cells with the inducible all-in-one PiggyBac plasmids containing inducible DjCas13d and selection with 1 μg/ml puromycin for 7 days to ensure plasmid integration. Differentiation procedures were then initiated and cells were induced for DjCas13d expression using Doxycycline (D3072, Sigma) at the middle time point of differentiation. 5-7 days after induction, flow analysis was performed to quantify the percent of cells expressing the effector in each experimental group using the GFP reporter. The GFP+ percentage of cells with targeting guide groups for each effector was normalized to that of the corresponding non-targeting guide group for evaluation of cell viability upon target RNA knockdown. Three biological replicates were performed.

### RNA-seq library preparation and sequencing

For HEK293FT cells, total RNA was extracted with the RNeasy Plus 96 Kit (Cat #74192, QIAGEN) 48h after transfection. For H1 cells, cell numbers were counted and normalized between different samples (different effectors, guides and replicates) 72h after transfection, and total RNA was extracted with the RNeasy Plus 96 Kit (Cat #74192, QIAGEN). Stranded mRNA libraries were prepared using the NEBNext II Ultra Directional RNA Library Prep Kit (NEB, Cat# E7760L) and NEBNext Poly(A) mRNA Magnetic Isolation Module (NEB, Cat #E7490). The libraries were sequenced on a partial NovaSeq lane with 150 nt paired end reads. ∼20M reads were demultiplexed per sample.

### RNA-seq analysis and pathway analysis of CasRx off targets

Sequencing reads were aligned to the hg38 Ensembl transcriptome using Kallisto (Bray et al., 2016). Mapping was carried out using default parameters except for a b value (number of bootstraps) of 100. Differential transcript expression was performed with Sleuth (Pimentel et al., 2017) using triplicates to compare between targeting and non-targeting conditions. Significantly differentially expressed transcripts were defined as having an adjusted p value < 0.1 and a beta value > 0.5. Volcano plots were generated in R using the package EnhancedVolcano (Blighe et al., 2019). Pathway analysis of CasRx off targets was performed using Enrichr (E. Y. Chen et al., 2013; Kuleshov et al., 2016; Xie et al., 2021) with the Molecular Signatures Database (MSigDB).

### RNA-seq Spike-In for total RNA quantification

To quantify total RNA amount accurately and determine if uniform transcriptome depletion has occurred following CasRx- or DjCas13-mediated transcriptome targeting, an equal amount of ERCC RNA Spike-In Mix (ThermoFisher, Cat #4456740) was added to the total RNA extracted from cell number-normalized H1 samples using the recommended dilution ratio before library preparation. After library preparation and NGS sequencing, the ratio of experimental reads to spike-in reads was calculated for all samples, and then normalized to the ratio of control samples (non-targeting guides) to get the total RNA amount relative to NT.

### RNA integrity analysis

To examine RNA integrity, electrophoresis was performed on the extracted RNA and the electrophoresis graphs were visualized on high sensitivity RNA chips using either Bioanalyzer (Agilent 2100 Bioanalyzer, G2939BA) (for experiments in HEK293FT) or TapeStation (Agilent 4200 TapeStation system, G2991AA) (for experiments in H1).

### Stem cell differentiation to NPC, HPC, neurons and RNA targeting experiments

For RNA targeting experiments in NPC and HPC, human embryonic stem cells (hESCs, H1 line, WiCell) were first transfected with inducible piggyBac-based all-in-one DjCas13d plasmids containing a puromycin resistance gene as mentioned above. For RNA targeting experiments in neurons, H1s were first transfected with inducible piggyBac-based all-in-one DjCas13d plasmids containing neomycin resistant gene by replacing the puromycin resistance gene in the piggyBac-based all-in-one DjCas13d plasmid with a neomycin resistance gene. After selection for plasmid integration with 1 μg/ml puromycin (NPC and HPC) or 100 μg/ml G418 Sulfate (neurons) for 7 days, differentiation procedures were performed as outlined below.

For differentiation to NPC, stem cells were passaged with Accutase (Innovative Cell Technologies) and plated at 30,000 cells per well into Matrigel-coated 96-well plates with N2B27 media (DMEM/F12 (Thermo Fisher) + N2 (100x, Thermo Fisher) + B27 without vitamin A (50x, Thermo Fisher)) containing ROCK inhibitor Y-27632 (10 uM, Abcam) and bFGF (40 ng/mL, Corning). The following day (day 0), media was replaced with N2B27 media containing AZD-4547 (50 nM, Abcam, Cat# ab216311), LDN-193189 (250 nM, Sigma, Cat# SML0559), A83-01 (250 nM, Sigma, Cat# SML0788), and XAV-939 (3 uM, Abcam, Cat# ab120897) to achieve dual SMAD and Wnt inhibition. Media was changed daily. On day 3, AZD-4547 was removed. On day 4, cells were passaged with Accutase (Innovative Cell Technologies) at 1:3 and plated again onto Matrigel-coated 96-well plates in N2B27 media containing ROCK inhibitor Y-27632 (10 uM, AbAcam), LDN-193189 (250 nM, Sigma, Cat# SML0559), A83-01 (250 nM, Sigma, Cat# SML0788), and XAV-939 (3 uM, Abcam, Cat# ab120897). Media was replaced the next day with N2B27 containing LDN-193189 (250 nM, Sigma, Cat# SML0559), A83-01 (250 nM, Sigma, Cat# SML0788), and XAV-939 (3 uM, Abcam, Cat# ab120897). Media was changed daily and cells were induced for DjCas13d expression using Doxycycline (D3072, Sigma) on day 5. On day 8, all drugs were removed and the media was changed with N2B27 only (DMEM/F12 + N2 (100x) + B27 without vitamin A (50x)). On day 10, the cells were assayed for target knockdown and NPC marker expression (Pax6 and Sox1) using flow cytometry.

For differentiation to HPC, stem cells were passaged with ReLeSR (StemCell Technologies) and plated at ∼40 colonies per well into Matrigel-coated 12-well plates with mTesR media (StemCell Technologies) containing ROCK inhibitor Y-27632 (10 uM, Abcam). The following day (day 0), media was replaced with 2 mL Hematopoietic Media A (STEMdiff Hematopoietic Basal Media (StemCell Technologies) with STEMdiff Hematopoietic Supplement A (200x, StemCell Technologies)). On day 2, a half-media change with Hematopoietic Media A was performed. On day 3, the media was fully replaced with 2 mL Hematopoietic Media B (STEMdiff Hematopoietic Basal Media (StemCell Technologies) + STEMdiff Hematopoietic Supplement B (200x, StemCell Technologies). On day 5, there was a half-media change with Hematopoietic Media B, and cells were induced for DjCas13d expression using Doxycycline (D3072, Sigma). On day 7 and day 10, 1 mL fresh Hematopoietic B media was added but no media was removed. On day 12, the cells were assayed for target knockdown and HPC marker expression (CD43) using flow cytometry.

For differentiation to neurons, hESCs (H1) were passaged with Accutase (Innovative Cell Technologies) and plated at 12,000 cells per well into Cultrex (R&D Systems 343400502)-coated 96-well plates with mTeSR media (StemCell Technologies) containing ROCK inhibitor Y-27632 (10 uM, Abcam). The following day cells were infected with lentivirus containing a doxycycline-inducible Ngn2 cassette in mTeSR media (StemCell Technologies) containing polybrene (10 mg/mL, Santa Cruz Biotechnology sc-134220). Following infection, media was changed daily to mTeSR media (StemCell Technologies). When cells reached 70% confluency, they were passaged with Accutase (Innovative Cell Technologies) and re-plated at 12,000 cells per well into Cultrex-coated 96-well plates with mTeSR media (StemCell Technologies) containing ROCK inhibitor Y-27632 (10 uM, Abcam). The day of passage was designated as day 0 of the differentiation protocol. The following day (day 1), media was replaced with mTeSR media (StemCell Technologies). On day 2, cells were induced for Ngn2 and DjCas13d expression using 2 ug/mL Doxycycline (2 ug/mL, Sigma D3072). On day 3, media was replaced with neural induction media (NIM, DMEM/F12 (Gibco 11330032) + Penicillin-Streptomycin (Gibco 15140122) + Doxycycline (2 ug/mL, Sigma D3072) + Laminin (1.2 ug/mL, Sigma L4544) + Insulin (5 ug/mL, Roche 11376497001) + BSA (10 mg/mL, Sigma A4161) + Apo-transferrin (10 mg/mL, Sigma T1147) + Putrescine (1.6 mg/mL, Sigma P57800) + Progesterone (0.00625 mg/mL, Sigma P8783) + Sodium selenite (0.00104 mg/mL, S5261) + BDNF (10 ug/mL, Sigma B3795) + Puromycin (10 ug/mL, Life Technologies A1113803)). Media was changed daily. After 3 days of puromycin selection, cells were passaged with Accumax (Innovative Cell Technologies) and plated at 87,500 cells per well with neural maturation media (Neurobasal differentiation media (Neurobasal Media (Gibco 21103049) + DMEM Media (Gibco 10569010) + HEPES (0.5x, Gibco 15630130) + Penicillin-Streptomycin (Gibco 15140122) + Glutamax (1 mM, Gibco 35050061)) + Doxycycline (2 ug/mL, Sigma D3072) + Laminin (2.4 ug/mL, Sigma L4544) + BDNF (10 ug/mL, Sigma B3795) + dbCAMP (49.14 ug/mL, Sigma Aldrich D0627) + B27 with vitamin A (1x, Gibco 17504044) + N-acetyl cysteine (5 ug/mL, Sigma A9165) containing ROCK inhibitor Y-27632 (10 uM, Abcam). Media was changed daily. On day 8, media was replaced with neural maturation media containing AraC (2.4 ug/mL, Sigma Aldrich C1768) to remove any post-mitotic neurons from the culture. On day 11, the cells were assayed for target knockdown using flow cytometry.

