## Supplementary Figures for "Deep learning and CRISPR-Cas13d ortholog discovery for optimized RNA targeting"

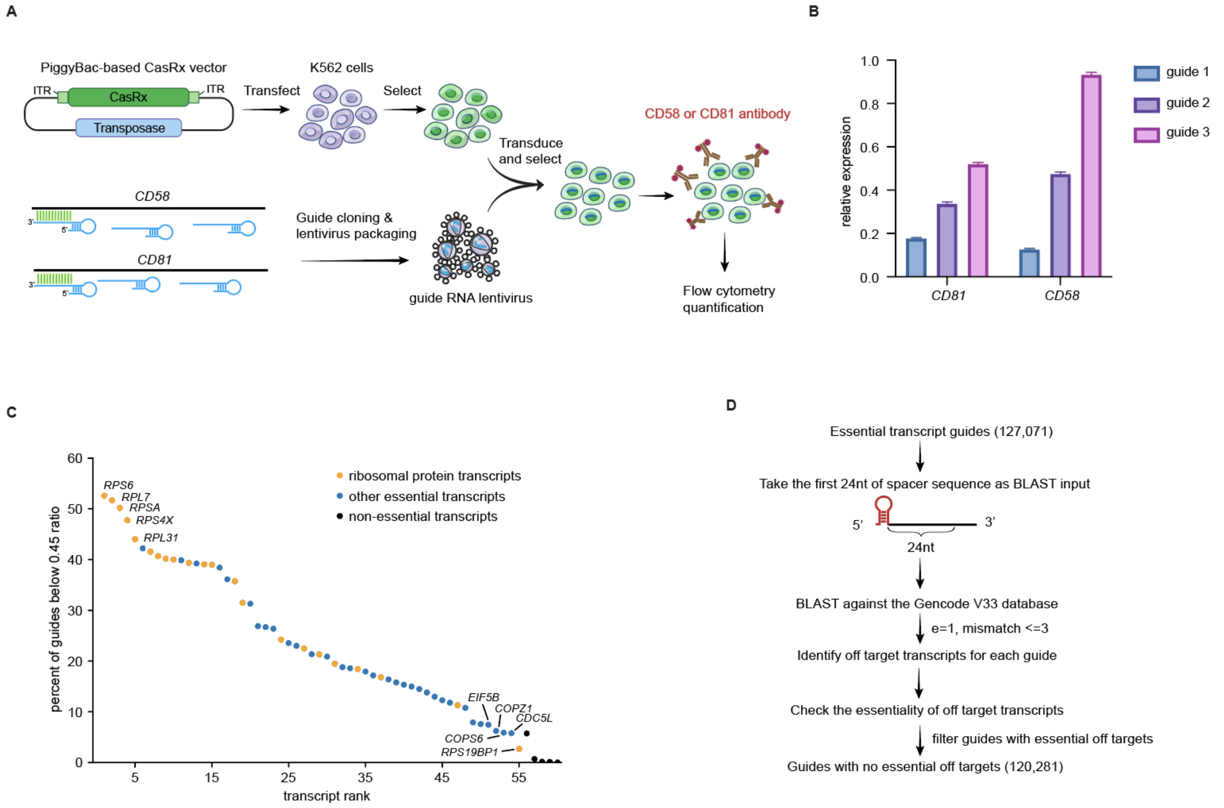


**Supplementary Figure 1: CasRx knockdown variability in K562 cells and survival screen data processing**

**A.** Schematic of single guide CasRx-mediated knockdown of transcripts encoding CD58 and CD81 in K562 cells with a flow cytometry readout. **B.** Comparison of cell surface protein expression of CD58 and CD81 in K562 cells following CasRx-mediated knockdown of the respective transcripts using three different guides per transcript, relative to non-targeting guide control. Mean ± SEM for n = 3 biological replicates. **C**. Transcript ranking based on the percentage of highly depleted guides from the survival screen depicted in **Figure 1A**. Individual transcripts are ranked based on the percentage of guides below the cut-off ratio (day14/input ratio <0.45). Orange dots denote ribosomal protein transcripts; blue dots denote other essential transcripts; black dots denote non-essential transcripts. The top 5 and bottom 5 essential transcripts are annotated. The full transcript list is provided in Table S2. **D.** Guide filtering approach based on off targets in other essential transcripts. The filtered dataset is provided in Table S4.

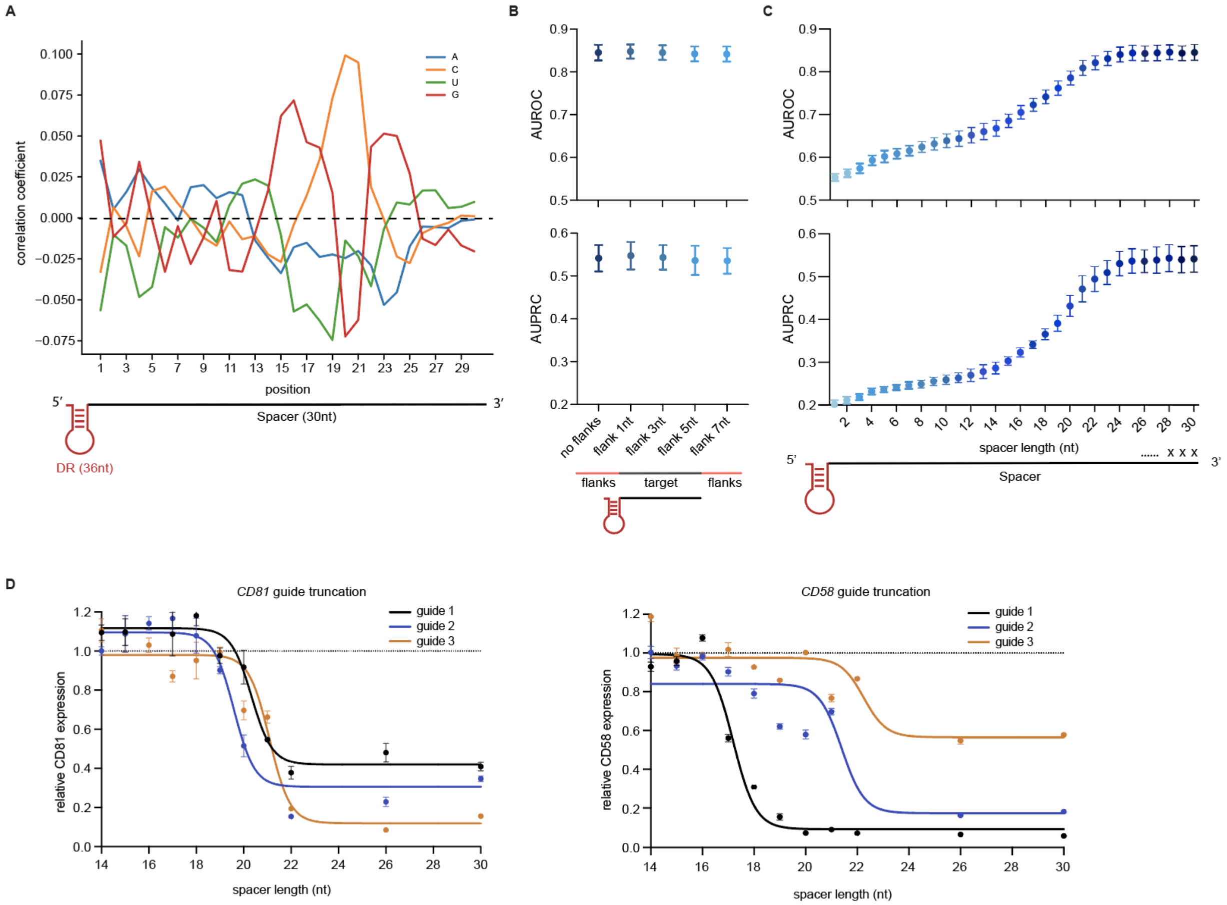


**Supplementary Figure 2: Contributions of CasRx guide sequence and length to guide efficiency. A.** Pearson correlation coefficient of each nucleotide with guide efficiency at each position along the 30 nt spacer. **B.** The effect of target RNA flanking sequence on model prediction accuracy. Target flanking sequences of various lengths (1, 3, 5, or 7 nt) were added to the 30 nt guide target sequence in the CNN model, and the resultant models were evaluated for guide efficiency prediction accuracy. Model AUROCs and AUPRCs (mean ± SD) are shown. **C.** The effect of guide spacer length on model prediction accuracy. The guide spacer sequence was computationally truncated in single-nucleotide intervals from the 3′ end, starting from the full-length 30 nt spacer sequence down to 1 nt. CNN models based on the truncated guide were evaluated for their performance, and AUROCs and AUPRCs (mean ± SD) are shown. **D.** Experimental validation of the knockdown efficiencies of the indicated guide spacer truncations when targeting *CD81* or *CD58*. Target protein knockdown in K562 cells was evaluated 48h after transfection by flow cytometry. Mean ± SEM for n = 3 replicates.

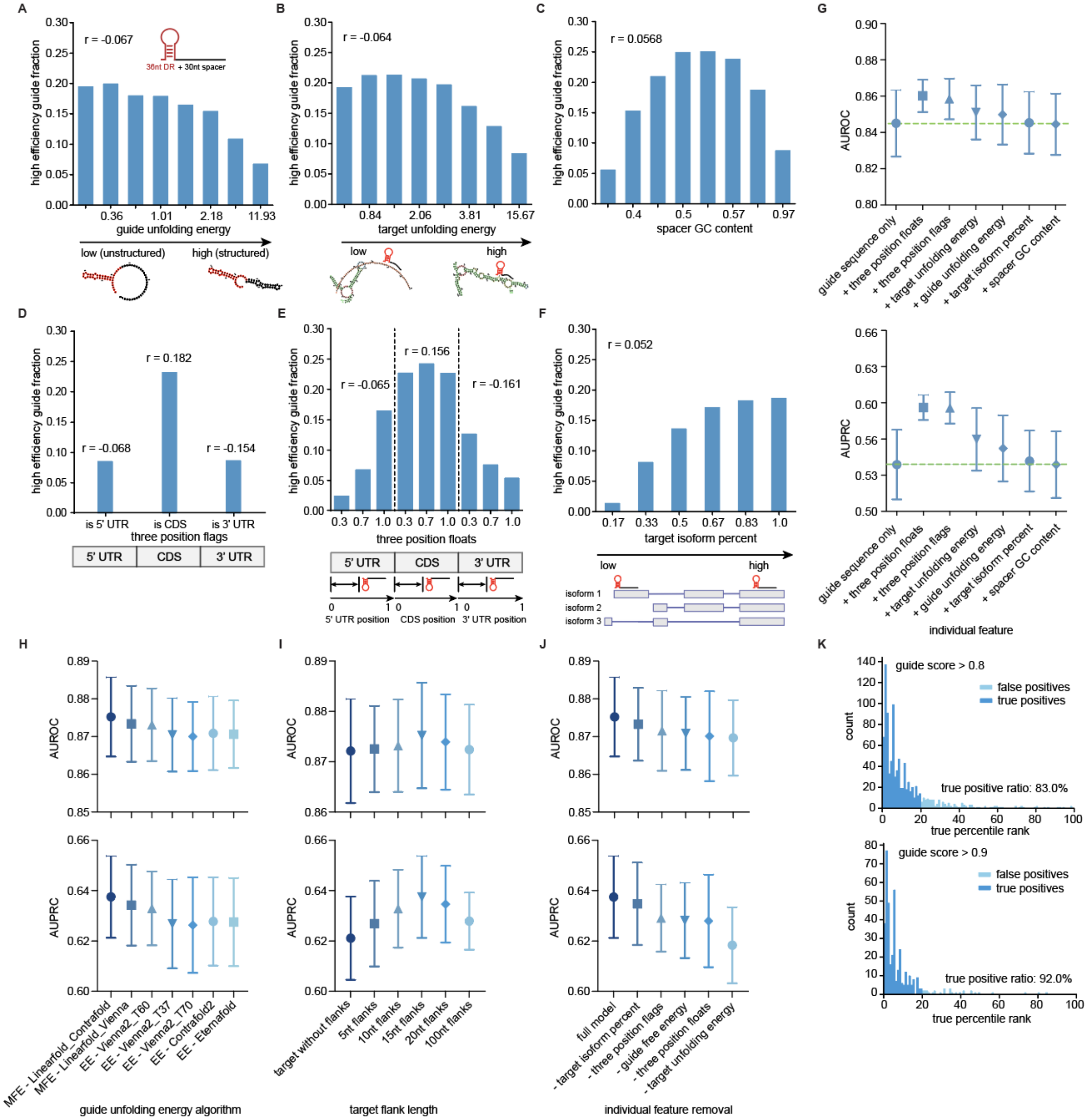

A
r = -0.067
36nt DR + 30nt spacer
high efficiency guide fraction
guide unfolding energy
low (unstructured)
high (structured)
B
r = -0.064
target unfolding energy
low
high
C
r = 0.0568
spacer GC content
D
r = 0.182
r = -0.068
r = -0.154
is 5' UTR
is CDS
is 3' UTR
three position flags
5' UTR
CDS
3' UTR
E
r = 0.156
r = -0.065
r = -0.161
three position floats
5' UTR position
CDS position
3' UTR position
F
r = 0.052
target isoform percent
low
high
isoform 1
isoform 2
isoform 3
G
AUROC
AUPRC
guide sequence only
+ three position floats
+ three position flags
+ target unfolding energy
+ guide unfolding energy
+ target isoform percent
+ spacer GC content
individual feature
H
MFE - Linearfold_Contrafold
MFE - Linearfold_Vienna
EE - Vienna2_T60
EE - Vienna2_T37
EE - Vienna2_T70
EE - Contrafold2
EE - Eternafold
guide unfolding energy algorithm
I
target without flanks
5nt flanks
10nt flanks
15nt flanks
20nt flanks
100nt flanks
target flank length
J
full model
- target isoform percent
- three position flags
- guide free energy
- three position floats
- target unfolding energy
individual feature removal
K
guide score > 0.8
false positives
true positives
true positive ratio: 83.0%
guide score > 0.9
true positive ratio: 92.0%
count
true percentile rank

**Supplementary Figure 3: Contributions of secondary features to guide efficiency prediction, feature optimization, and final model performance**

**A.** Fraction of high efficiency guides across different predicted guide RNA unfolding energies based on the LinearFold algorithm (Huang et al., 2019). Higher guide unfolding energy corresponds to a more structured guide RNA. **B.** Fraction of high efficiency guides across different predicted target unfolding energies based on the LinearFold algorithm. Higher target unfolding energies correspond to a more structured and less accessible target site. **C.** Fraction of high efficiency guides belonging to bins representing different spacer GC content. **D.** Fraction of high efficiency guides across target positions binned by 5′ UTR, coding sequence (CDS) or 3′ UTR. **E.** The 5' UTR, CDS, and 3' UTR regions were each divided positionally into 3 bins, and the fraction of high efficiency guides located within each target position bin are plotted. **F.** Target RNA positions were classified by their level of conservation across all known isoforms of the target transcript, calculated using the Refseq database. The fraction of high efficiency guides targeting sites falling into each of the 6 bins of target isoform conservation is shown. **G.** Model performance following addition of individual secondary features. Each secondary feature (or feature group) was added to the sequence-only CNN model individually, and model performance was evaluated by AUROC and AUPRC (mean ± SD) on held-out transcripts in the 9-fold split. Features are ordered based on the magnitude of their effect on final model performance. Green dashed lines denote the AUROC and AUPRC for the guide sequence only model. **H.** Comparison of model performance incorporating predicted RNA folding energies using different RNA secondary structure algorithms. The guide MFE (minimum free energy) was calculated using either the CONTRAfold or Vienna model with the LinearFold algorithm. The ensemble unfolding energy (EE) was calculated using Contrafold2, Eternafold, and Vienna2. Model AUROC and AUPRC (mean ± SD) are shown for each algorithm. **I.** Model performance upon adjustment of target flank length for target RNA unfolding energy calculation. Bidirectional target flanks with different lengths (0, 5, 10, 15, 20 or 100 nt) were added to the 30 nt target site to calculate the local target unfolding energy. Model AUROC and AUPRC (mean ± SD) are shown for each target flank length. **J.** Model performance upon removal of individual secondary features. Each secondary feature (or feature group) was removed individually from the final CNN model and the resultant model AUROC and AUPRC (mean ± SD) are shown. Removal of each secondary feature reduced model accuracy, supporting their necessity in the final model. **K.** Distribution of the true percentile rank of guides above different model score thresholds. The model returns a score ranging from 0 to 1 for every guide. The true percentile rank distributions of guides above two model score thresholds (0.8, upper plot and 0.9, lower plot) are shown (1st percentile = most efficient guides, 99th percentile = least efficient guides). True positives (top 20th percentile) are plotted in dark blue, and false positives (20th-100th percentile) are plotted in light blue.

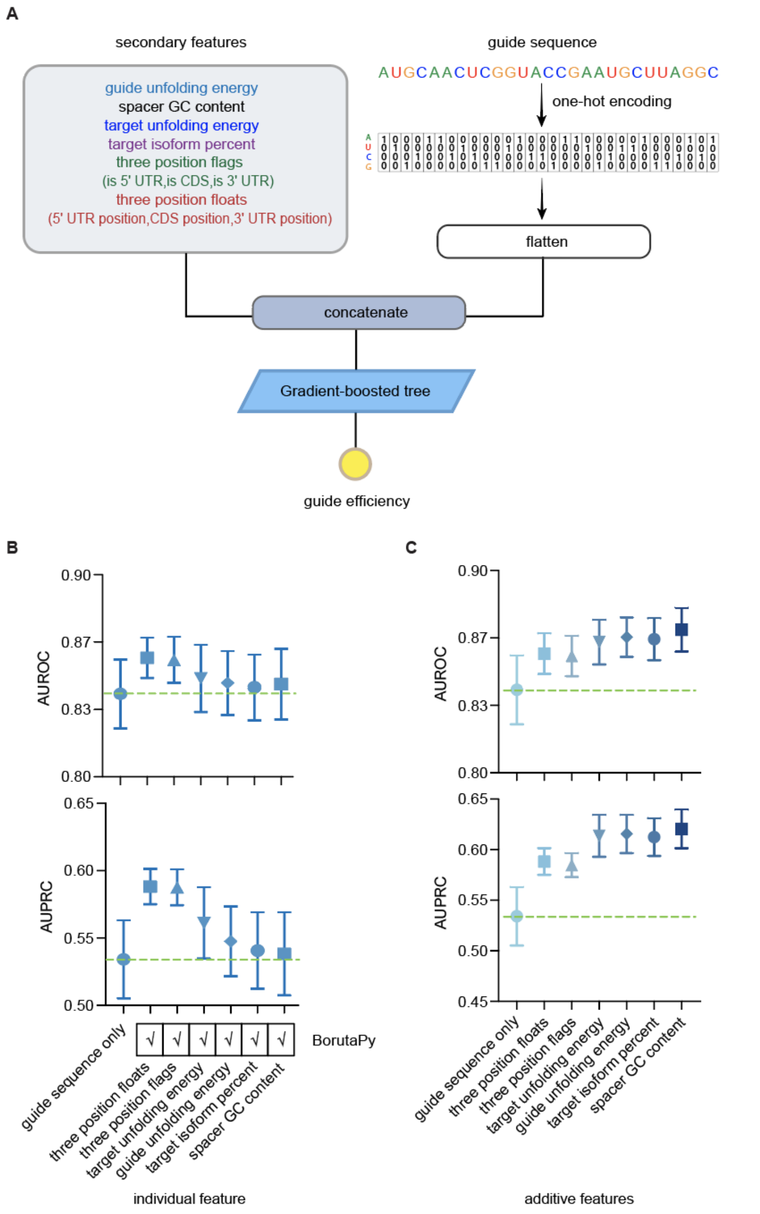


**Supplementary Figure 4: Addition of secondary features also improves the gradient-boosted tree (GBT) model. A.** Schematic of secondary feature addition to the GBT model – the best performing model not based on deep learning. **B.** GBT model performance, evaluated by AUROC and AUPRC for held-out transcripts across all 9 fold test-training data splits, upon addition of individual secondary features. Features were sorted based on final model performance. The table below the plots indicates the necessity of each feature in the model, as evaluated using BorutaPy (Kursa et al., 2010). Green dashed lines denote the average AUROC and AUPRC for the guide sequence only model. **C.** GBT model performance upon sequential cumulative feature addition, ordered by each feature's individual contribution to model performance from panel B. Model AUROC and AUPRC (mean ± SD) are shown. Overall, the final GBT model performed slightly worse than the CNN model (shown in **Fig. 1F**).

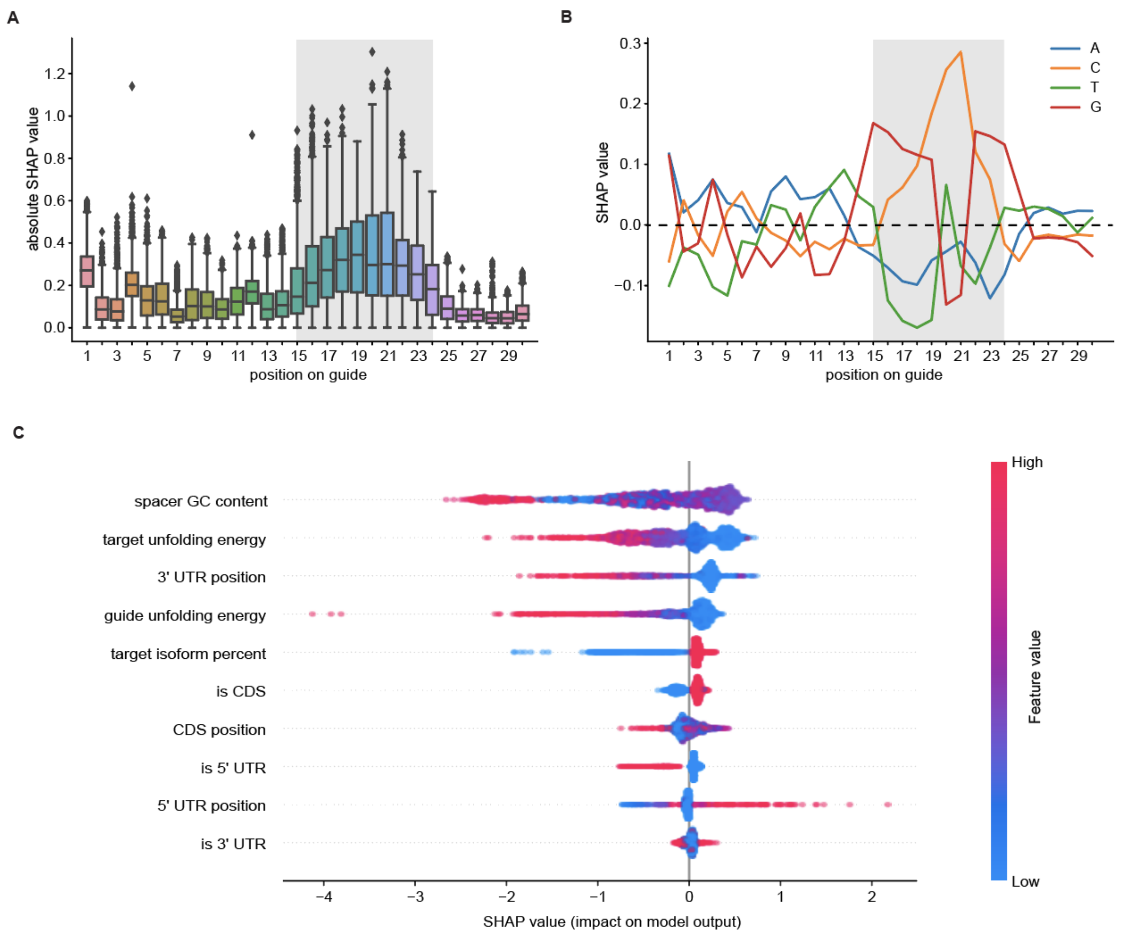


**Supplementary Figure 5: Model interpretation of the GBT model using SHapley Additive exPlanations (SHAP). A.** Evaluation of the importance of each position in the guide spacer sequence in the GBT model using SHAP (Lundberg et al., 2020), with the core region (nt 15-24) highlighted in gray shading. **B.** Evaluation of the importance of each positional nucleotide in the guide sequence in the GBT model using SHAP values. The gray shaded box denotes the core region of the spacer sequence. **C.** Contribution of secondary features to guide efficiency prediction in the GBT model. The beeswarm plot displays the SHAP values (impact on model output) against feature input values for all secondary features across all test guides. Positive feature input values are indicated in red while negative feature input values are indicated in blue. The features are ranked based on the sum of their absolute SHAP values across all test set guides.

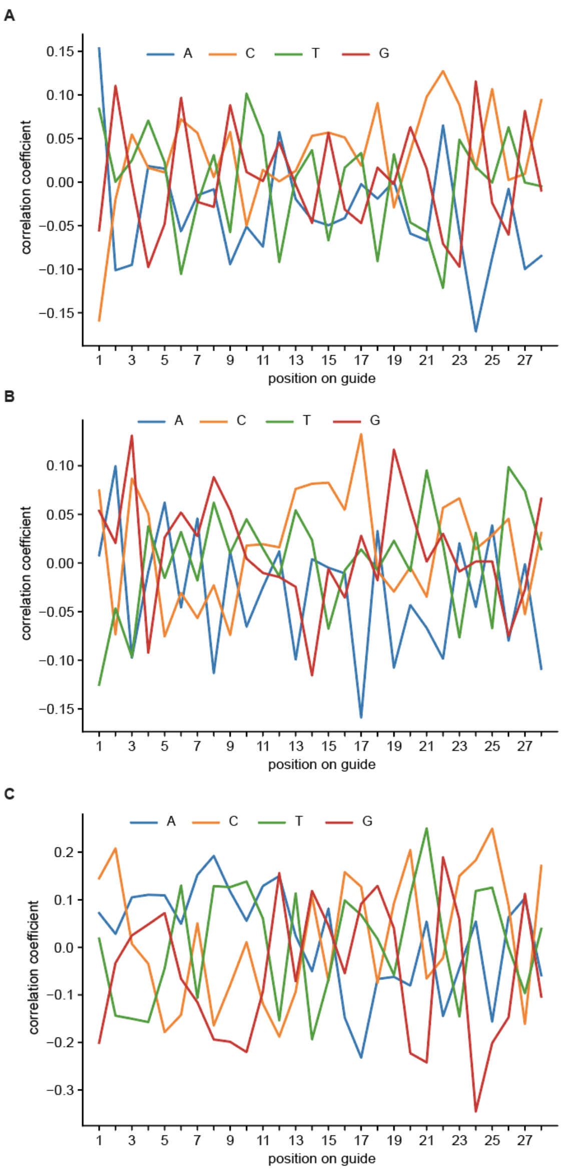


**Supplementary Figure 6: Absence of consistent Cas13a guide sequence correlation with guide efficiency across published datasets. A.** Correlation of each nucleotide with guide efficiency at each guide position in the LwaCas13a luciferase knockdown dataset (Abudayyeh et al., 2017; Metsky et al., 2021). 186 LwaCas13a guides for *Gluc* and 93 guides for *Cluc* were analyzed and the Pearson correlation coefficient for each positional nucleotide with guide efficiency is shown. **B.** Correlation of each nucleotide with guide efficiency at each guide position in the LwaCas13a endogenous transcript knockdown dataset (Abudayyeh et al., 2017; Metsky et al., 2021). 279 LwaCas13a guides for *KRAS, PPIB* and *MALAT1* were analyzed and the Pearson correlation coefficient for each positional nucleotide with guide efficiency is shown. **C.** Correlation of each nucleotide with guide efficiency at each guide position in the LwaCas13a ADAPT dataset (Metsky et al., 2022). 85 perfect match guides from LwaCas13a ADAPT data were analyzed and the Pearson correlation coefficient for each positional nucleotide with guide efficiency is shown.

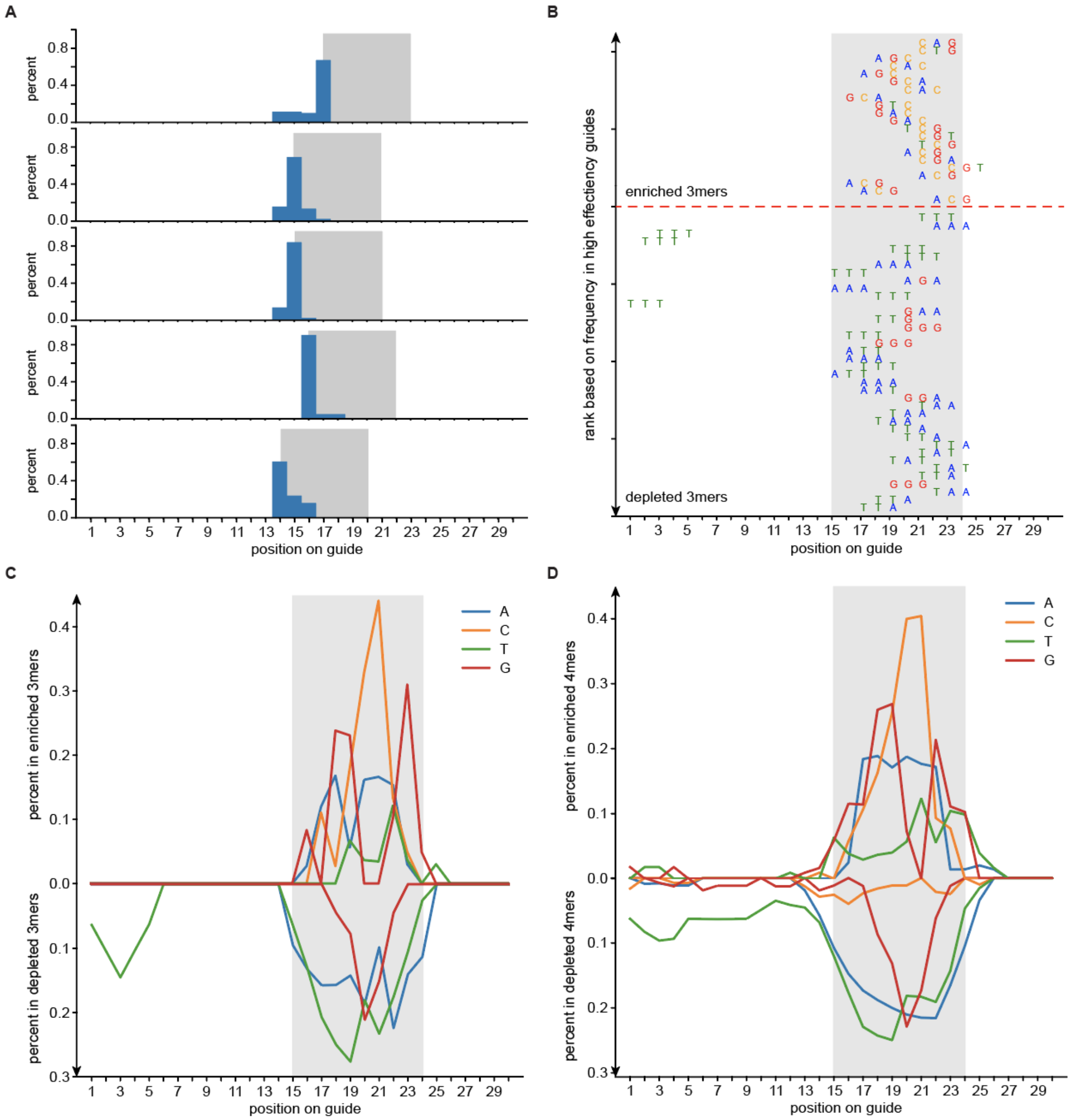


**Supplementary Figure 7: Investigation of favored sequence motifs for high efficiency guides. A.** The positional distribution of the seqlets (sequence regions with high importance based on IG scores) along the 30 nt spacer in the top 5 patterns identified by TF-MoDISco. The histogram summarizes the start position distribution of the seqlets in each pattern. The gray box highlights the pattern window starting at the mode position in each pattern. The identified patterns are highly positional. **B.** Top enriched and depleted 3-mers at each position, ranked by their frequency in high efficiency guides. The percentage of all possible 3-mers at each position in high efficiency guides and non-high efficiency guides was calculated, and enriched (or depleted) 3-mers were selected based on their enrichment (or depletion) ratio in high efficiency guides to non-high efficiency guides. **C.** Summary of base composition from top enriched and depleted 3-mers at each position in high efficiency guides. **D.** Summary of base composition from enriched and depleted 4-mers at each position in high efficiency guides.

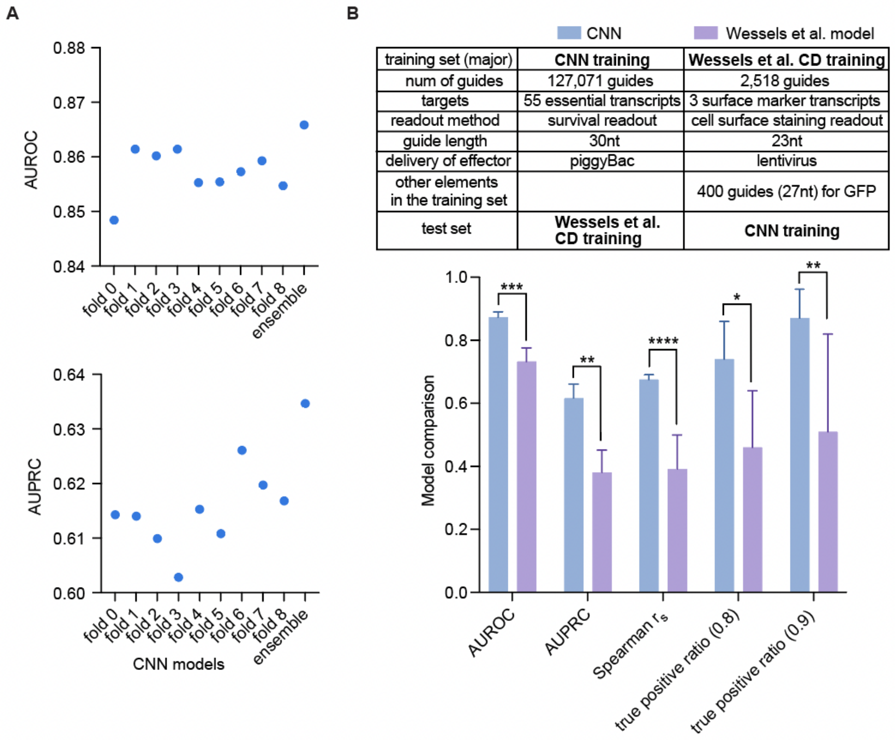


**Supplementary Figure 8: Improved performance of our deep learning model based on head-to-head comparison with a previously reported model. A.** Comparison of model performance of individual CNN models from the 9-fold split of the survival screen data relative to the ensemble CNN model which averages the prediction of individual models. Model AUROC and AUPRC on the two validation targets (*CD58* and *CD81*) are shown. **B.** Comparison of the previously published CasRx random forest model (Wessels et al., 2020) to the CNN model described here on the opposing dataset. The top 20% efficient guides for each transcript are defined as high efficiency guides for model evaluation. Model AUROC, AUPRC, Spearman's correlation coefficient and true positive ratio at 0.8 and 0.9 model score cutoffs (mean ± SD) across transcripts are shown. * P<0.05, ** P<0.01, *** P<0.001, **** P <0.0001 based on Welch’s t test.

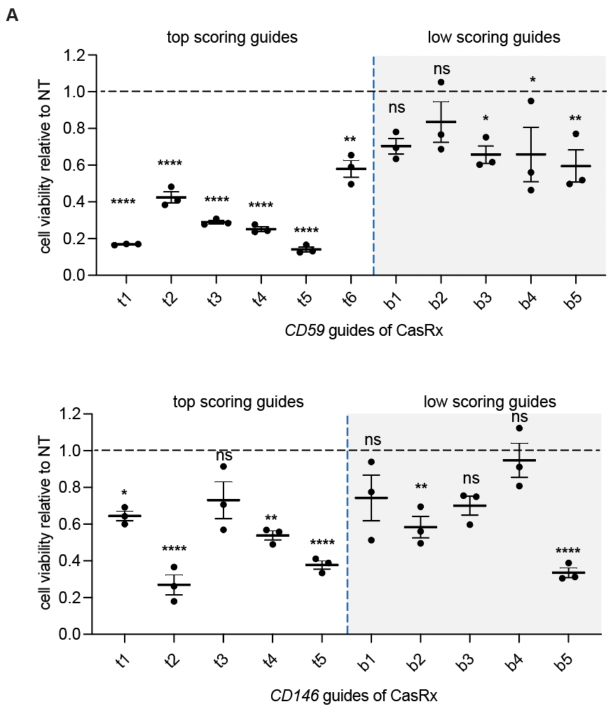


**Supplementary Figure 9: Observed cellular toxicity of CasRx in A375 cells. A.** Cellular viability of A375 cells with CasRx targeting *CD59* and *CD146*. Mean ± SEM for n = 3 replicates for 5 or 6 top scoring guides and 5 low scoring guides (same as in Fig. 3G) is shown. P values are based on ordinary one-way ANOVA for each guide compared to NT guide controls.

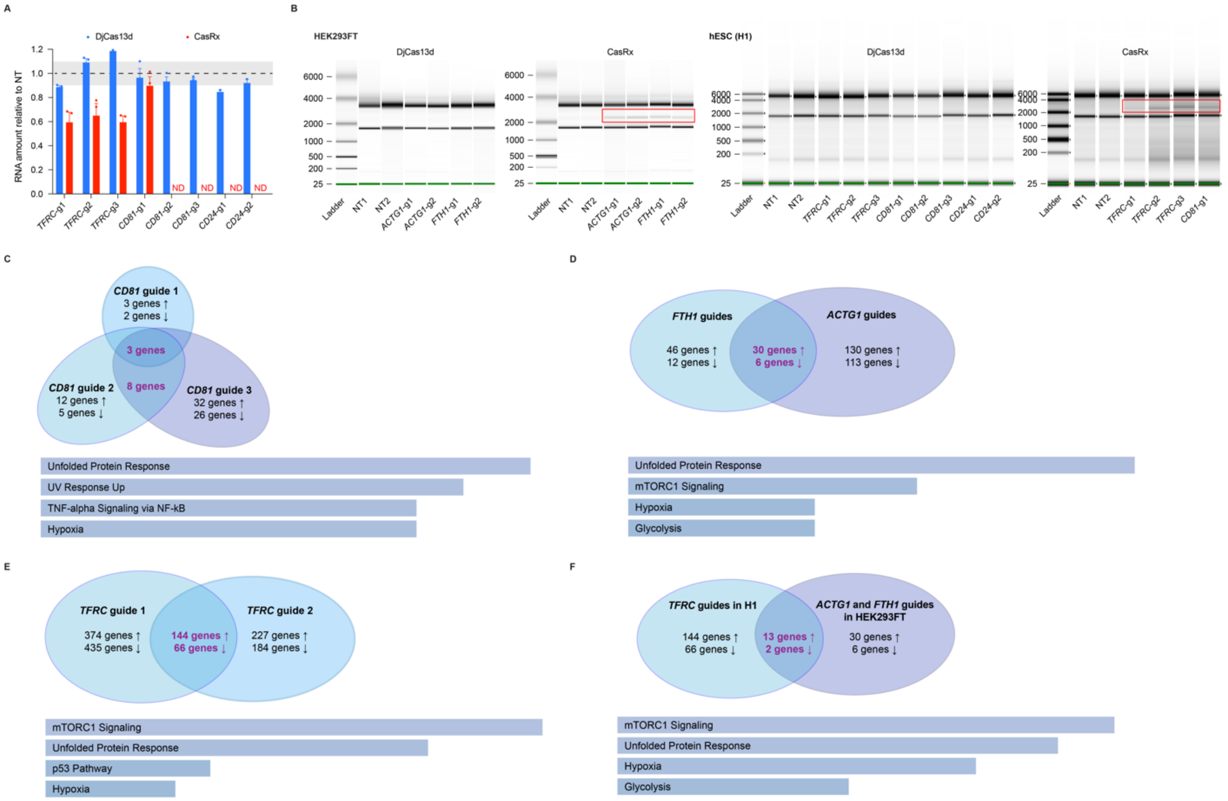
A
DjCas13d
CasRx
RNA amount relative to NT
1.2
1.0
0.8
0.6
0.4
0.2
0.0
ND
ND
ND
ND
TFRC-g1
TFRC-g2
TFRC-g3
CD81-g1
CD81-g2
CD81-g3
CD24-g1
CD24-g2
B
HEK293FT
DjCas13d
CasRx
6000
4000
2000
1000
500
200
25
Ladder
NT1
NT2
ACTG1-g1
ACTG1-g2
FTH1-g1
FTH1-g2
hESC (H1)
DjCas13d
CasRx
TFRC-g1
TFRC-g2
TFRC-g3
CD81-g1
CD81-g2
CD81-g3
CD24-g1
CD24-g2
C
CD81 guide 1
3 genes ↑
2 genes ↓
3 genes
CD81 guide 2
12 genes ↑
5 genes ↓
8 genes
CD81 guide 3
32 genes ↑
26 genes ↓
Unfolded Protein Response
UV Response Up
TNF-alpha Signaling via NF-kB
Hypoxia
D
FTH1 guides
46 genes ↑
12 genes ↓
30 genes ↑
6 genes ↓
ACTG1 guides
130 genes ↑
113 genes ↓
Unfolded Protein Response
mTORC1 Signaling
Hypoxia
Glycolysis
E
TFRC guide 1
374 genes ↑
435 genes ↓
144 genes ↑
66 genes ↓
TFRC guide 2
227 genes ↑
184 genes ↓
mTORC1 Signaling
Unfolded Protein Response
p53 Pathway
Hypoxia
F
TFRC guides in H1
144 genes ↑
66 genes ↓
13 genes ↑
2 genes ↓
ACTG1 and FTH1 guides in HEK293FT
30 genes ↑
6 genes ↓
mTORC1 Signaling
Unfolded Protein Response
Hypoxia
Glycolysis

**Supplementary Figure 10: Transcriptome-wide RNA integrity and specificity in cells treated with CasRx and DjCas13d**

**A.** Comparison of total RNA amount extracted from cells expressing CasRx or DjCas13d with the indicated guides, normalized to cell number and relative to a non-targeting guide using exogenous RNA spike-ins in hESCs (H1). The black dashed line indicates relative RNA amount at 1.0 (average of NT guides) and the shaded box indicates the SEM for NT guides. Mean ± SEM for n = 3 replicates. ND, no data because of the low number of cells collected owing to CasRx's toxicity in these conditions. DjCas13d's targeting guides showed no significant RNA amount differences from NT guides while CasRx's *TFRC*-targeting guides had significantly lower (P <0.0001) RNA amount compared to NT guides. * P<0.05, ** P<0.01, *** P<0.001, **** P <0.0001 based on ordinary one-way ANOVA compared to NT guide conditions. **B**. Electrophoresis graphs of total RNA from CasRx and DjCas13d guides in HEK293FT (left panel) and H1 (right panel). The red box indicates a band that appears only in CasRx-treated cells with targeting guides. Sharp, clear bands around 5 kb and 1.9 kb represent the 28S and 18S rRNAs, respectively, and the absence of additional bands or smears indicates intact RNA. Degraded RNA shows additional bands or appears as a lower molecular weight smear. **C.** Overlap of the differentially affected transcripts by CasRx among the three *CD81*-targeting guides in HEK293FT cells. Pathway analysis of overlapping genes reveals enriched pathways (based on MSigDB Hallmark 2020) ranked by p value (E. Y. Chen et al., 2013; Kuleshov et al., 2016; Xie et al., 2021). The top 4 pathways are shown, with the length of the bars representing the significance of the pathway. **D.** Overlapping off-targets by CasRx among the *ACTG1* and *FTH1* targeting guides in HEK293FT cells. Pathway analysis of overlapping genes reveals enriched pathways (based on MSigDB Hallmark 2020) ranked by p value (Chen et al., 2013; Kuleshov et al., 2016; Xie et al., 2021). The top 4 pathways are shown, with the length of the bars representing the significance of the pathway. **E.** Overlap of the differentially affected transcripts by CasRx among the two *TFRC*-targeting guides in H1. Pathway analysis of overlapping genes reveals enriched pathways (based on MSigDB Hallmark 2020) ranked by p value (Chen et al., 2013; Kuleshov et al., 2016; Xie et al., 2021). The top 4 pathways are shown, with the length of the bars representing the significance of the pathway. **F.** Overlap of the differentially affected transcripts by CasRx between *TFRC*- targeting guides in stem cells (H1) and *ACTG1*- and *FTH1*- targeting guides in HEK293FT cells. Pathway analysis of overlapping genes reveals enriched pathways (based on MSigDB Hallmark 2020) ranked by p value (Chen et al., 2013; Kuleshov et al., 2016; Xie et al., 2021). The top 4 pathways are shown, with the length of the bars representing the significance of the pathway.

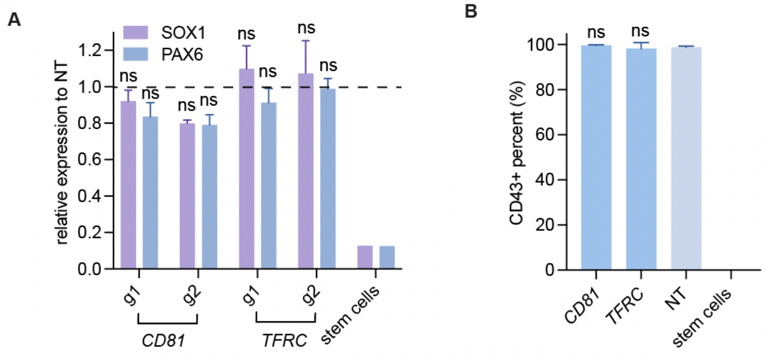


**Supplementary Figure 11: DjCas13d has no negative impact on differentiation efficiency when targeting various transcripts in stem cell-derived NPCs and HPCs**

**A.** Relative expression level of NPC markers (SOX1 and PAX6) as measured by flow cytometry in stem cell-derived NPCs following DjCas13d-mediated targeting of *CD81* and *TFRC* with the indicated guides, relative to a non-targeting guide control. The expression level of these marker genes in the originating stem cell line is shown on the right for comparison. Mean ± SEM for n = 3 replicates. **B.** Percent of stem cell-derived HPCs expressing the cell-surface HPC marker CD43 following DjCas13d-mediated targeting of *CD81* or *TFRC,* as measured by flow cytometry. The percentage of HPCs expressing CD43 in the non-targeting guide condition (NT) and the percentage of originating stem cells expressing CD43 are shown as controls. Mean ± SEM for n = 3 replicates.
